# Signatures in domesticated beet genomes pointing at genes under selection in a sucrose-storing root crop

**DOI:** 10.1101/2024.05.27.596033

**Authors:** Amar Singh Dhiman, Demetris Taliadoros, Eva H. Stukenbrock, J. Mitchell McGrath, Nazgol Emrani, Christian Jung

## Abstract

The genus *Beta* encompasses important crops such as sugar, table, fodder, and leaf beets. All cultivated beets are believed to have originated from the wild sea beet, *B*. *vulgaris* subsp. *maritima*. Sugar beet, a recent crop dating back nearly 200 years, was selectively bred for enhanced root yield in combination with high sucrose content. We assembled a *Beta* diversity panel comprising wild and cultivated beet accessions. Whole-genome sequencing identified 10.3 million SNP markers. Four distinct genetic clusters were identified: table beet, sugar beet, Mediterranean sea beet, and Atlantic sea beet. A phylogenetic analysis revealed that cultivated beet accessions were genetically closer to Mediterranean than to Atlantic sea beet and that cultivated beets producing storage roots share a common ancestor. Cultivated beets exhibited genome regions with reduced nucleotide diversity compared to Mediterranean sea beets, indicating selection signatures. These regions contained putative candidate genes with potential roles in root development, suppression of lateral root formation, flowering time, and sucrose metabolism. A yet unknown sucrose transporter on chromosome 6 showed reduced nucleotide diversity exclusively in sugar beet accessions compared to other *Beta* types with low sucrose content, suggesting its role in sucrose storage. Within a region of high nucleotide diversity between accessions with contrasting root phenotypes, we found two genes encoding auxin response factors, which play a crucial role in root development. We reason these genes to be significant root thickening regulators in root crops.

## Introduction

The order Caryophyllales (family Amaranthaceae) contributes important crops such as spinach, quinoa, amaranth, and buckwheat. Within this order, the genus *Beta* comprises cultivated and wild forms. Sugar beet, fodder beet (syn. mangelwurzel), leaf beet (syn. mangold, Swiss chard, chard, spinach beet), and table beet (syn. garden beet, red beet, beet root) belong to the species *Beta vulgaris* ssp. *vulgaris*. They are closely related to their wild relatives, *B. vulgaris* ssp. *maritima*, *B*. *vulgaris* ssp. *adanensis, B. patula,* and *B. macrocarpa*. The cultivated lineages show pronounced phenotypic differences owing to their usage. It is believed that cultivated beets originated from the wild progenitors of *B*. *vulgaris*. ssp. *maritima*, also known as ‘sea beet’ ^1^.

Leaves of the Mediterranean sea beet populations have been consumed as a vegetable for over 2000 years. The enlarged root types (historically termed ‘Runkelrüben’) have been speculated to originate in the Near East (Iraq, Iran, and Turkey) and then spread west ^2^. However, the period of domestication is still uncertain.

Table and leaf beet are important vegetables cultivated in temperate and tropical climates across the globe. The petioles of leaf beets exhibit a rich spectrum of colors, encompassing shades of white, yellow, pink, and red. Table beets are primarily grown for their succulent roots and hypocotyls, which exhibit a wide range of diversity in terms of shape, including round, globe-shaped, flattened, and cylindrical roots. The red color in their roots is attributed to the presence of betalain pigments, while yellow hues result from betaxanthin pigments. The accumulation of betalains in beets is regulated by two loci, namely *R* and *Y*, which have been identified and molecularly described. The *BvCYP76AD1* gene at the *R* locus encodes a cytochrome P450 enzyme essential for betalain biosynthesis ^3^. Conversely, the *Y* locus contains the anthocyanin MYB-like gene *BvMYB1*, determining the presence or absence of pigmentation (red or yellow) in the flesh of beet roots ^4^. Notably, the dried red beet powder, known for its antioxidant properties, is employed as a natural food dye in the food industry ^5, 6^.

In 1747, the German chemist A.S. Marggraf found that ‘Runkelrüben’ roots store sucrose (then known familiarly as ‘cane sugar’). This incentivized growing Runkelrüben for sucrose production by the end of the 18^th^ century. The sucrose content of beet roots at that time was around 4% (fresh weight). Selection for root dry mass started after the first sugar factory was built in Silesia in 1801, then a part of Germany. Mass selection to improve open-pollinated populations was the major breeding method for the first hundred years before introducing progeny testing ^7^. This resulted in an immense increase in root weight and sucrose content ^8, 9^. Today, sugar beets are cultivated in more than 50 countries around Europe, Asia, the Americas, and parts of Africa (FAOSTAT, 2022: OECD/FAO (2022). Most modern beet varieties are F_1_ hybrids. In Europe, root yield ranges between 60 and 85 t/hectare, and sucrose content ranges between 17 and 19% fresh weight. In contrast, the sucrose concentration in fodder beet is lower, ranging between 4 and 10%^10^.

Wild sea beets produce small, non-spherical spangled roots characterized by numerous lateral/secondary branches emerging from a single taproot ^5^. Remarkably, there are no discernible phenotypic differences in the roots of leaf beets compared to sea beets. However, table beets, fodder beets, and sugar beets exhibit significantly enlarged storage organs, predominantly formed by a non-branched primary taproot and, to some extent, by the hypocotyl.

Sea beet roots possess supernumerary root cambia, although they are not notably swollen compared to the roots of table, fodder, and sugar beets. It can be hypothesized that the presence of supernumerary root cambia might have facilitated the selection of roots with swollen characteristics. Nevertheless, there is considerable variation in cambium ring numbers in both wild and cultivated beets. Typically, the roots of fodder and table beets have 3-5 cambium rings, and sugar beets can produce up to 12 successive concentric rings of cambia ^9^. Table and fodder beets produce an enlarged hypocotyl, constituting a substantial part of the storage organ, in contrast to sugar beets, where the root is the primary storage organ ^1^. Cultivated beets with thick storage roots have been intensively selected against lateral or secondary root branching, resulting in a thick single taproot. The development of thick storage roots can be attributed to concurrent and synchronous increases in cell division and enlargement within these cambial rings ^5, 11^.

Cultivated beets are biennial species that form a large leaf and root mass in the first year. The floral transition occurs after winter, initiating with shoot elongation termed bolting and the production of up to 10,000 florets ^9^. Flowering must be avoided entirely during field production due to a drastic reduction in sucrose yield. Beets are an allogamous crop species due to a complex gametophytic self-incompatibility in wild and cultivated forms, preventing self-pollination. However, self-incompatibility is overcome by a self-fertility locus with a self-fertility allele *SF*, allowing inbred line production for hybrid breeding ^1^.

Sea beets are abundant around the Mediterranean and at the cost of Western and Northern Europe. Although they belong to the same species, their phenological development and morphology are strikingly different from cultivated root types. They remain a vital genetic resource for various biotic stress resistances^12^.

Cultivated beets and wild forms are diploid (2n=2x=18) with an estimated genome size ranging from 714 to 758 Mb ^13^. The beet genome is highly complex and repetitive, with approximately ∼63% of its content comprising repetitive elements ^14^. The first reference genome assembly was derived from the doubled-haploid sugar beet ‘KWS2320’ and comprised ∼567 Mbp, with 85% of the sequences assigned to 9 chromosomes and predicting 27,421 protein-coding genes ^15^. The inbred line ‘EL10’ genome was recently published using long-read sequencing, BioNano optical mapping, DoveTail Hi-C scaffolding, and Illumina short-reads. The ‘EL10.2_2’ genome assembly comprises ∼569 Mb, with 99.2% of the sequences assigned to 9 chromosomes and 24,186 predicted protein-coding genes ^16^. Two more fully annotated *de novo* genome assemblies have been published for one *B*. *vulgaris* ssp. *maritima* (‘WB42’) and one *B*. *patula* accession (‘BETA548’) ^17, 18^. The assembled genome sizes for *B*. *vulgaris* ssp. *maritima* and *B*. *patula* were 590 Mb and 624 Mb, respectively.

Whole-genome sequencing studies have evaluated genetic diversity within and between cultivated and wild beets. Galewski and McGrath (2020) analysed genetic diversity and phylogenetic relationships among 23 cultivated beet accessions, representing a broad phenotypic variation for foliar and root traits. Each accession was treated as a population due to the pooled sequencing of individual plants belonging to that particular accession. Table beets were differentiated as a distinct crop type compared to sugar beets. Fodder and chard accessions were intermediate between table and sugar beet ^8^. Another study estimated phylogenetic relationships among 606 sugar beet and wild beet accessions using reference-free distance-based approaches^19^. Sea beets clustered into Mediterranean and Atlantic accessions. Sea beets from Greece were closest to sugar beet, which made the authors suggest that sugar beet might originate from this region. In a recent study, genomic data from 656 *Beta* section 1 genomes were compared to characterize genes undergoing selection during the domestication and breeding between wild and cultivated accessions ^20^. In total, 10 million variants were identified across the 656 beet accessions, encompassing wild and cultivated forms. Across the sugar beet genomes, various regions totaling 15 Mb were characterized under artificial selection due to low genetic variation. These variation-poor regions were enriched for genes involved in shoot system development, stress response, and carbohydrate metabolism.

In our study, a worldwide collection of wild and cultivated accessions was grown in the field to assess agronomically important phenotypes. After whole-genome sequencing, the population structure of cultivated beet lineages and their wild relatives was assessed. We depicted genome regions with remarkably high sequence diversity between lineages with contrasting phenotypes such as root thickening and bolting. Low nucleotide diversity within cultivated lineages highlighted regions under intense selective pressure during the past 200 years. Candidate genes were localized within these regions with putative functions in storage root formation, sucrose storage, and seed morphology. These genes are proposed to be key genes for root crop domestication and sucrose storage.

## Materials and Methods

### Plant materials and growth conditions

First, we selected 485 wild and cultivated beet accessions from the Plant Breeding Genebank at CAU, Kiel, representing diverse geographical origins. Seeds were visually inspected to characterize the seed types (monogerm vs. multigerm), and germination rates were assessed in the greenhouse. Plants were grown in row plots in the field in Kiel (Germany) between May and October 2020 to assess the phenological development, phenotypic characters, and homogeneity within accessions. Based on these data, 290 phenotypically homogenous wild and cultivated beet accessions were chosen as a *Beta* mini-core collection for further analyses (Supplementary Table 1) (Figure 1))

**Figure 1:**
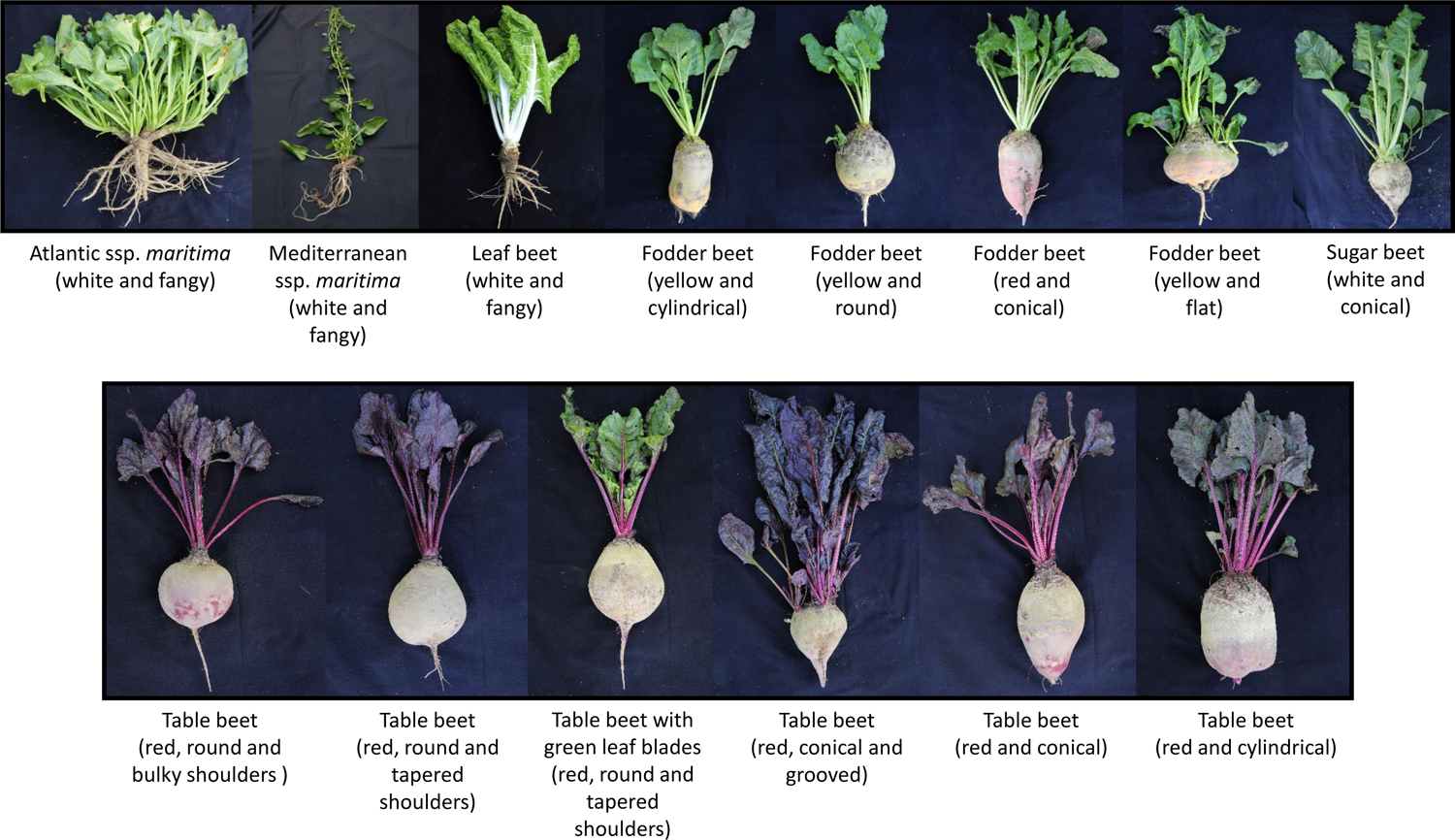
Phenotypic diversity displayed among wild and cultivated beets captured in the study. The upper panel displays, from left to right, a non-bolting Atlantic sea beet plant with white, fangy, and branched root; a bolting Mediterranean sea beet plant with white, fangy, and branched root; a leaf beet plant with broad light green wrinkled leaves, a white and depressed petiole, and sea beet-like fangy root; a fodder beet plant with yellow and cylindrical thickened storage root; a fodder beet plant with yellow and round thickened storage root; a fodder beet plant with orange-red external root color and presenting sugar beet-like conical thickened storage roots with a prominent groove; a fodder beet plant with yellow-orange and flat thickened storage root; and a sugar beet plant with a thickened white conical storage root, a source of sucrose. The lower panel, from left to right, illustrates a table beet plant with dark red leaves, petioles, and a round, thickened storage root with bulky shoulders; a table beet plant with dark red leaves, petioles, and a round, thickened storage root with tapered shoulders; a table beet plant with green leaf blades and red veins, red-pink petioles, and a round, thickened storage root with tapered shoulders; a table beet plant with dark red leaves, petioles, and a sugar beet-like conical thickened storage root with a prominent groove; a table beet plant with dark red leaves, petioles, and a sugar beet-like conical thickened storage root without a groove also known as ‘smooth root’; a table beet plant with dark red leaves, petioles, and a cylindrical thickened storage root; and top view of cross-sections of different table beet roots displaying a target-like appearance due to the accumulation of betalain pigments in the parenchyma and/or cambium storage root tissues.

The mini-core collection was grown in a field near Göttingen (Germany) in 2021 and 2022. At the end of April, seeds were hand-sown as single-row plots with seven plants/row in a randomized complete block design with three replications. Each replication block contained 290 row plots, each comprising a single accession. Each row plot contained seven plants per accession. Each replication block was divided into two sub-blocks. Each sub-block was 13.5m × 8m in size. The distances between rows and between plants were set to 45 cm and 20 cm, respectively. In total, 870 row plots were sown in all three replication blocks. Individual plants from each accession were phenotyped for external root color and root shape, leaf and petiole color, and bolting habit (Figure 1).

### DNA extraction and whole-genome sequencing

Leaf samples were collected from young leaves of six-week-old single plants grown in the field. Leaf samples were transported on dry ice and were immediately lyophilized. For DNA extraction, one representative plant was selected from each accession at the end of the cultivation season 2021. DNA was isolated using the QIAGEN DNeasy plant mini kit following their standard protocol. The purity and quality of DNA were assessed by agarose gel electrophoresis, and the concentration was determined using a Nano Drop2000 spectrophotometer (ThermoFisher Scientific, Waltham, United States).

Novogene (United Kingdom) performed whole-genome sequencing using the short reads on the Illumina NovaSeq 6000 sequencing platform. We aimed to achieve an average of 12 Gb of paired-end (PE) 2 x 150 bp reads with quality Q>30 Phred score per sample. This coverage is approximately equivalent to ∼15x of the haploid sugar beet genome (∼758 Mb).

### Raw reads quality check and alignment

Raw sequence reads were obtained in the zipped FASTQ format. Quality check for the raw reads was performed using the FastQC v0.11.9 ^21^ and MultiQC v1.13 ^22^ algorithms. Raw reads were trimmed, and library bar-code adapters were removed with trimmomatic-v0.39 ^23^ using the following criteria: ILLUMINACLIP:adapters.fa-:2:30:10, LEADING:3; TRAILING:3; SLIDINGWINDOW:4:15; MINLEN:36. Filtered reads were used for the further downstream analysis. A long-read genome assembly of a sugar beet inbred line EL10 was used as a reference genome ^16^. Filtered reads were aligned to the EL10.2_2 reference genome assembly (Phytozome genome ID: 782), using BWA-MEM (v-0.7.17) ^24^ with default parameters. The sequence alignment files (SAM) from each accession were sorted and indexed individually using SAMtools ^25^. Read grouping was assigned using AddOrReplaceReadGroups, and PCR duplicates were marked using MarkDuplicates on each bam file using Picard tools (http://broadinstitute.github.io/picard/).

GATK best practices were followed to call and filter the variants ^26^. Firstly, variants were identified with GATK (v4.2.5.0) ^27^ using HaplotypeCaller function on each bam file in a genomic variant call format (GVCF) mode. Then, individual g.vcf files from 290 samples were consolidated into a single GVCF file using CombineGVCFs function of GATK. Finally, SNPs and small InDels were identified with a joint calling approach using the GenotypeGVCFs function of GATK. To obtain high confidence variants, SNPs were hard-filtered using parameters “QDL<L2.0 | | FSL>L60.0 | | MQL<L40.0 | | SORL>L3.0 | | MQRankSumL<L−12.5 | | ReadPosRankSumL<L−8.0”, and indels with “QDL<L2.0 | | FSL>L200.0 | | SORL>L10.0L| | MQRankSumL<L−12.5 | | ReadPosRankSumL<L−8.0”. The hard-filtered variants with genotype calls with a depth < 2 and > 50 were excluded. Finally, only bi-allelic variants were kept, and variants with a missing rate of >10% or a minor allele frequency (MAF) of <0.05 were removed, which resulted in a high confidence (HC) set of ∼11.7 million variants, including SNPs and small InDels. The complete bioinformatics pipeline is shown as a workflow (Supplementary Figure 1).

### Variant annotation

We annotated the HC variant set using the Ensemble Variant Effect Predictor (VEP) ^28^ based on the EL10.2_2 sugar beet genome and annotation (Phytozome genome ID:782). We used VEP to categorize variants in coding regions based on their features: synonymous, missense, splice acceptor, splice donor, splice region, start lost, start gained, stop lost, and spot retained. Gene content, SNP density, and INDEL density within 100 kb non-overlapping windows were plotted using Circos ^29^. Then, we used the HC variant set for phylogeny, population stratification, demography, and selective sweep analyses.

### Phylogeny and population structure analyses

For clustering analysis, the variants were pruned for linkage-disequilibrium (LD) to reduce the bias and redundancy. The independent variants, i.e., LD pruned variants, were used for PCA (Principal Component Analysis) and admixture analysis. Therefore, the HC variant set was pruned for LD with PLINK (version 1.9) ^30^ using a window size of 10 kb with a step size of one SNP and an r2 threshold of 0.5, resulting in a 1.4 million pruned variants. A PCA was performed on LD pruned variants using the SNPrelate package in R ^31^. We estimated the top 10 principal components. The first (PC1) and second (PC2) principal components were plotted using custom R scripts. A neighbor-joining (NJ) tree was constructed using PHYLIP with 100 bootstraps. The p-distance matrix was generated from the variant call format (VCF) file using VCF2Dis (v1.47), fneighbor, and fconsense (Version: EMBOSS:6.6.0.0 PHYLIPNEW:3.69.650). An NJ tree was constructed with 100 bootstraps using PHYLIP (version 3.696), consensus was generated, and the tree layout was generated using FigTree. The accession with the lowest breadth of coverage and furthest placement on PC1 from the rest of the accessions was used to root the tree.

A population structure analysis was performed using ADMIXTURE (Version: 1.3.0) ^32^. We analyzed population structure with cluster K ranging from 1–10 using a default fivefold cross-validation (--cvL=L5). Each K was run with ten replicates. Obtained Q matrices were aligned using Pong ^33^. Highly admixed sugar beet, table beet, and Mediterranean sea beet accessions were discounted, and only accessions with >60% genetic ancestry threshold at *K*=4 were chosen for further analyses.

### Linkage disequilibrium analysis

The LD was calculated for each subpopulation or genetic cluster using the HC variants set. Then, LD was calculated for the whole population, including wild accessions. The LD was calculated on SNP pairs within a 500 kb window using default parameters in PopLDdecay (v 3.31) ^34^. The LD decay was plotted using custom R scripts based on the ggplot2 package. The LD decay was measured as the distance at which Pearson’s correlation efficiency (*r*^2^) dropped to half of the maximum.

### Population genomics analysis

Pairwise population differentiation (*F_ST_*) and nucleotide diversity (π) were calculated. Individuals were divided into subpopulations based on the clustering analysis. Nucleotide diversity (π) for subpopulations and genetic differentiation (FST) between different subpopulations were performed within a non-overlapping 10 kb sliding window and 1 kb steps using VCFtools (version 0.1.13) ^35^. *F_ST_* values were calculated between wild and cultivated beet populations using the 10 kb non-overlapping window approach. The top 5% ratios of mean *F_ST_*values between wild and cultivated population regions were considered candidate regions for population divergence.

The inbreeding coefficient (F) was calculated using VCFtools (version 0.1.13) ^35^ to evaluate heterozygosity at variant loci in individual accessions. The inbreeding coefficient across different cultivated lineages was plotted using custom R scripts.

### Identification of domestication sweeps

Selective sweeps in the sugar beet reference genome were identified using three different approaches, calculating XP-CLR ^36^ scores with the parameters ‘-w 1 0.005 200 1000 - p0 0.95’, *F*_ST_, and nucleotide diversity (π) ratios, using VCFtools (v0.1.13) ^35^. All three approaches involve the comparison of allele frequencies within and between subpopulations. We extracted the top 5% of scores/values from XP-CLR and *F*_ST_ analyses. The overlapping regions between both analyses were considered as potential regions under selection. To consolidate these overlapping regions, we used the bedtools merge option. We then identified the genes located within these regions by using bedtools intersect with the .GFF file.

We applied nucleotide diversity (π) as a criterion to further refine our candidate selective sweep regions. Specifically, we focused on regions with exceptionally low nucleotide diversity within cultivated lineages but with high nucleotide diversity within *B. vulgaris* ssp. *maritima* accessions. Within these regions exhibiting low genetic diversity, we gave special attention to genes related to sucrose transport, inositol transport, and auxin metabolism.

### Demography history of beet evolution

MSMC2 (version 2.1.4) ^37^ was used to estimate the effective population size. Briefly, VCFtools was used to generate population-wise VCF files without any missing data. Using default parameters, the SNPs within each sub-population were phased using BEAGLE (version 5.4) (Browning and Browning 2007). Eight randomly chosen samples from each subpopulation were selected from the population VCF file. Then, a masked file was generated using bamCaller.py for each sample. Then, the ‘generate_multihetsep.py’ script within MSMC tools was used to create input files for MSMC2 for each chromosome separately. MSMC2 was then run on the phased SNPs with a generation time 1 and the mutation rate per generation per site as 4L×L10^−8^.

## Results

### Assembling a *Beta* core collection

Based on the passport data, we selected 485 *B*. *vulgaris* L. accessions with diverse geographical origins, including Atlantic and Mediterranean sea beet accessions, to capture the extensive genetic diversity in wild sea and cultivated beets. Together, these accessions encompass the entire phenotypic variation of the species.

First, an observation trial was performed in the field, and accessions displaying phenotypic heterogeneity or not corresponding to the seed bank’s passport data were discarded. Next, we curated the passport data, ensuring only phenotypically true-to-type accessions were included in a *Beta* mini-core collection assembly. The remaining heterogeneous accessions were discarded. As a result, a *Beta* mini-core collection was assembled, consisting of 290 highly homogeneous accessions encompassing both wild sea beets and cultivated beets. It incorporates germplasm from various sources, including the USDA (Fort Collins and East Lansing, USA), the IPK Genebank (Gatersleben, Germany), and collections from the Plant Breeding Institute, Kiel, as well as commercial hybrids from Germany. This collection includes both annual and biennial accessions from *Beta vulgaris* ssp. *vulgaris* and *B*. *vulgaris* ssp. *maritima*. The mini-core collection comprises 91 *B*. *vulgaris* ssp. *maritima*, 81 sugar beet, 59 table beet, 30 fodder beet, and 29 leaf beet accessions from 40 countries worldwide. This collection represents the large phenotypic diversity of this genus regarding phenological development, storage root formation, sucrose storage, and leaf and root coloration (Figure 1).

### Whole genome sequencing of a *Beta* core collection reveals high genetic variation among different crop types

Whole genome sequencing of 290 accessions was performed, and about 4.2 Tb of whole-genome sequencing data was generated after sequencing a representative plant from each accession. In total, 28.1 billion raw reads were generated (Supplementary Table 2). After trimming the adapter content, 27.6 billion reads were retained (Supplementary Table 3). The accessions’ GC content (%) ranged between 35% to 38%. The coverage depth ranged between 15.4× and 45.75× per sample, with an average depth of ∼19.86× per sample (Supplementary Table 4).

Filtered reads from all accessions, including *B*. *vulgaris* ssp. *maritima*, were mapped to the long-range sugar beet genome assembly EL10.2_2. The average mapping rate of single reads to this genome was 98.61%, and the average mapping rate for both read pairs correctly mapped to the reference was 87.36%. In general, accessions belonging to cultivated lineages showed greater mapped reads and properly paired mapped reads than wild accessions. Sugar beet accessions exhibited the highest percentage of total mapped reads, ranging from 97.73% to 98.94%, and both pairs properly mapped reads, ranging from 85.65% to 89.45%. The *B*. *vulgaris* ssp. *maritima* total mapped reads ranged from 97.62% to 98.75%, and both-pairs-properly-mapped-reads ranged from 84.17% to 88.05%.

The mean coverage breadth achieved was 85.62% of the reference EL10.2_2 genome assembly, with sugar beet accessions exhibiting the highest average coverage fraction of 86.55% (83.72% to 88.72%). As expected, wild sea beet accessions showed a lower average coverage fraction of 84.91% (80.28% to 87.21%) (Table 1, Supplementary Table 4).

**Table 1:** Summary statistics of mapped reads from 290 beet accessions using the EL10 genome as a reference.

| Population | Number of accessions | Total mapped reads (%) |  |  | Both pairs properly mapped reads (%) |  |  | EL10 genome covered (%) |  |  |
| --- | --- | --- | --- | --- | --- | --- | --- | --- | --- | --- |
|  |  | Max. | Min. | Average | Max. | Min. | Average | Max. | Min. | Average |
| Sugar beet | 81 | 98.94 | 97.73 | 98.71 | 89.45 | 85.65 | 88.21 | 88.7 | 83.72 | 86.55 |
| Table beet | 59 | 98.83 | 98.29 | 98.67 | 88.48 | 86.67 | 87.74 | 87.83 | 82.95 | 85.52 |
| Fodder beet | 30 | 98.76 | 98.29 | 98.62 | 88.34 | 86.56 | 87.55 | 87.72 | 83.64 | 85.78 |
| Leaf beet | 29 | 98.73 | 97.36 | 98.57 | 88.3 | 85.92 | 87.09 | 87.03 | 82.89 | 85.32 |
| Mediterranean sea beet | 69 | 98.75 | 97.62 | 98.51 | 88.05 | 85.31 | 86.61 | 87.21 | 81.71 | 85.19 |
| Atlantic sea beet | 22 | 98.7 | 98.14 | 98.41 | 86.75 | 84.17 | 85.7 | 86.71 | 80.28 | 84.03 |

Approximately 85 million variant sites, including single-nucleotide polymorphisms (SNPs) and insertions/deletions (INDELs) were genotyped across 290 accessions with reference to the EL10.2_2 genome assembly. We filtered for biallelic variants with a maximum of 10% missing data, a minor allele frequency >5%, and a mean depth >5 across all genotyped accessions. After filtering, 11,669,554 high confidence (HC) variants were identified across 290 accessions. Of these, 10,365,662 (88.8%) were SNPs, and 1,303,892 (11.2%) were small INDELs (Supplementary Table 5).

The variant density was high across the genome, averaging 20.5 variants/kb. Generally, the highest variant density (SNPs and INDELs) was observed at the chromosomal ends. The average density of SNPs across 290 accessions was 1822 SNPs/100kb, ranging from 0 to 4659 SNPs/100kb. The average number of INDELs across 290 accessions was 229.2 INDELs/100kb, ranging from 0 to 601 INDELs/100kb (Figure 2). The number of variants across different chromosomes ranged from 1.1 to 1.5 million (Supplementary Table 5, Supplementary Figure 2), with an average of 1.29 million/chromosome. The highest number of variants were observed on chromosome 5, regardless of chromosome length.

**Figure 2:**
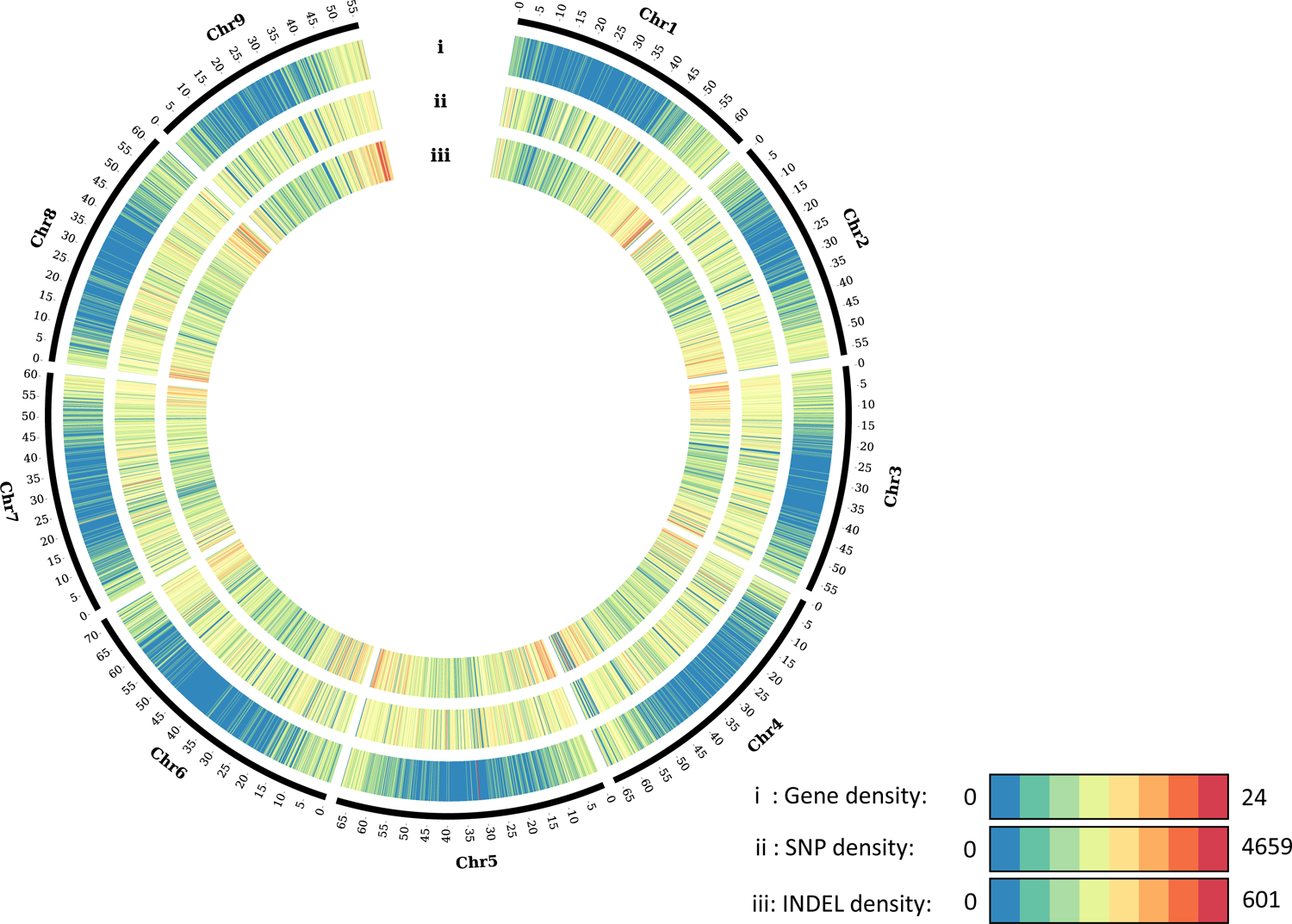
Visualization of 11 million variants (SNPs and INDELs) of the Beta mini-core collection across nine EL10 sugar beet chromosomes. The CIRCOS plot shows three tracks (outside to inside) representing (i) gene density, (ii) SNP density, and (iii) INDEL density across nine sugar beet chromosomes. Chromosome lengths are in Mb. Gene, SNP, and INDEL densities were calculated in 100 kb non-overlapping windows.

For SNP variant positions, 3,019,720 (29.2 %) SNPs were located in upstream and downstream regions of a predicted gene, 1,865,832 (18.0 %) SNPs were located within predicted intronic regions, and 4,845,240 (46.8 %) SNPs were located within intergenic regions (Supplementary Table 6). Only 479,462 (4.6 %) SNPs were within predicted open reading frames. Of these, only 211,493 SNPs (44.1 %) were predicted to be within a coding region, with 267,969 (55.9 %) being synonymous variants and 206,807 (43.1 %) being missense variants. Only 3,481 (0.73 %) of the variants led to ‘stop gained’ or ‘non-sense’ mutations (Supplementary Table 7).

Of INDEL variants, 736,158 (56.5 %) were deletions, and the remaining 566,978 (43.5 %) were insertions concerning the reference genome. In total, 448,030 (34.4 %) INDELs were located in upstream and downstream regions of a predicted gene, 304,470 (23.4 %) were located within predicted introns, and 504,179 (38.7 %) of INDELs were located in intergenic regions (Supplementary Table 6). Only 16,213 (1.2 %) INDELs were within predicted open reading frames. For INDELs altering the predicted polypeptide sequence, 9,252 (57.1 %) were predicted as frameshift mutations (Supplementary Table 7).

### Population structure of the beet diversity panel

Leaf beet accessions were grouped with the cluster of the Mediterranean Sea beet accessions. PC2, which accounted for 3.1 % of the variance, primarily separated the different cultivated beet lineages (Figure 3A, Supplementary Figure 3). We excluded the sea beet accessions from a second PCA analysis to further discern the genetic sub-clusters within the cultivated beet lineages. In this case, PC1 explained 4.4 % of the variance and effectively differentiated table beet accessions from sugar beet accessions, with each cultivated lineage forming a cloud of its distinct cluster along PC1.

**Figure 3:**
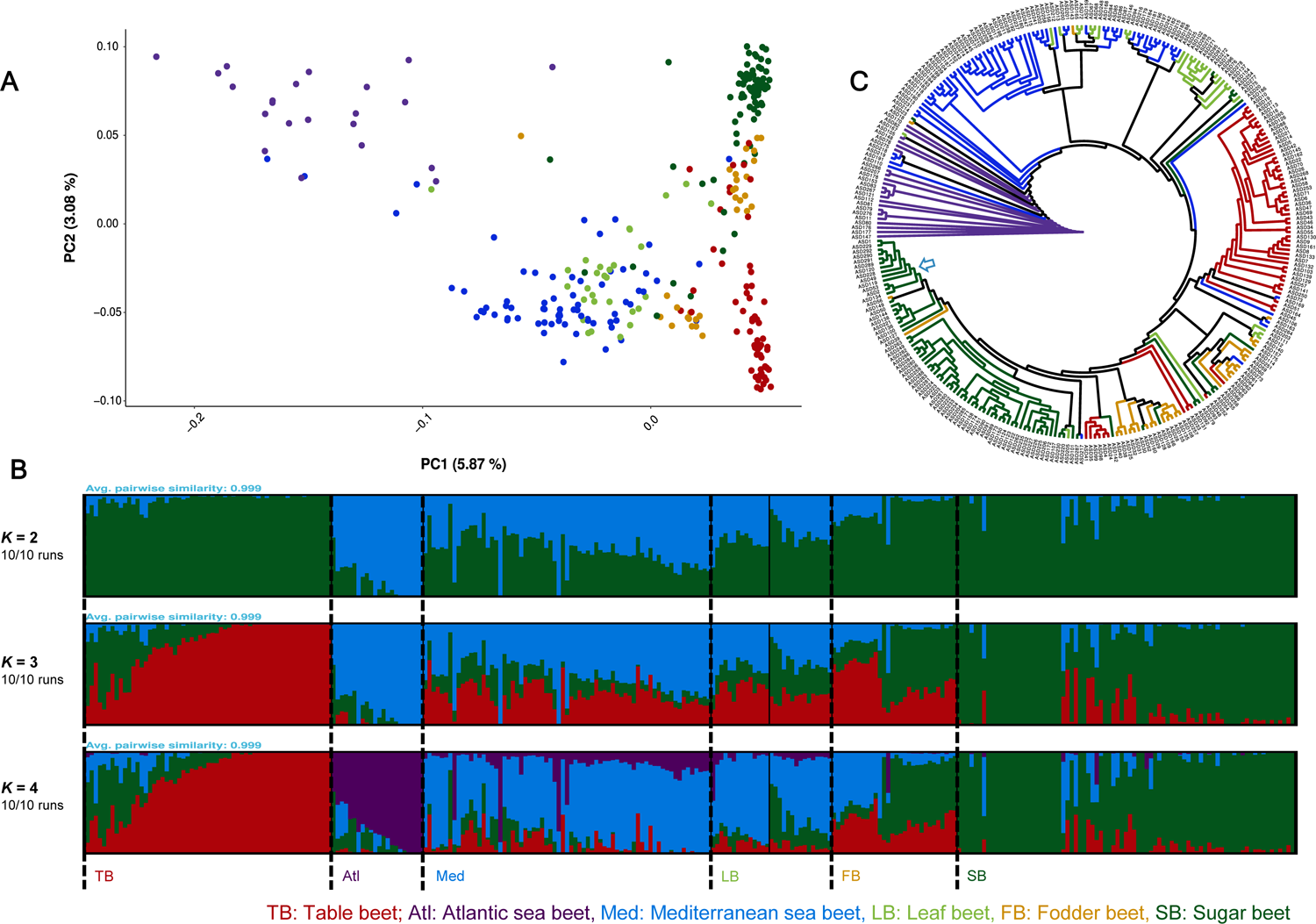
Genetic diversity, population structure, and Neighbor-joining phylogenetic tree of 290 wild and cultivated beet accessions from the Beta mini-core collection. (A) Principal component analysis (PCA) of 290 beet accessions. PC1 and PC2 represent the first two analysis components, accounting for 5.87% and 3.08% of the total variation, respectively. The colors represent different genetic clusters according to their beet type, representing the wild accession from the Atlantic (purple) and the Mediterranean (blue), sugar beet (dark green), table beet (red), fodder beet (orange), and leaf beet (light green) accessions. (B) The population structure analysis was conducted on the Beta mini-core collection using model-based clustering in ADMIXTURE, with different numbers of ancestral kinships (K=2 and K=10). K=4 was determined as the optimal number of populations. The graphs illustrate the ancestry proportions at K=2, 3, and 4. The Y-axis represents the share of genetic ancestry, while the X-axis represents different accessions. In the plot, blue represents the Mediterranean sea beet ancestry, green represents sugar beet ancestry, purple represents the Atlantic sea beet ancestry, and red represents table beet ancestry. (C) Neighbor-joining phylogenetic tree of the 290 accessions. Based on PCA and the breadth of coverage, the tree was rooted using an Atlantic sea beet accession as an outgroup. The color of the branches corresponds to the color codes used in PCA. A blue arrow indicates hybrid accessions from private breeding companies. An orange arrow indicates the fodder, table, and sugar beet accessions forming conical roots.

Fodder beet and leaf beet accessions showed less pronounced divergence and appeared intermediate between sugar beet and table beet accessions. Additionally, we observed a distinct separation between annual and biennial leaf beet accessions (Supplementary Figure 4).

As a next step, we performed a population structure analysis with the ADMIXTURE software. We used cross-validation errors to estimate the most suitable number of populations. At *K* = 2, two distinct clusters of *B*. *vulgaris* ssp. *maritima* and *B*. *vulgaris* ssp. *vulgaris* were formed. However, table beet and sugar beet accessions, in this case, shared a similar genetic ancestry. At *K*=3, a distinct cluster of table beet accessions was evident. At *K*=4, minimum cross-validation was observed (Supplementary Figure 5), with four distinct genetic clusters (Figure 3B) corresponding to table beet, sugar beet, Mediterranean sea beet, and Atlantic sea beet ancestries. Two distinct genetic clusters of wild sea beet accessions with different ancestry were evident, as well as two distinct genetic clusters within *B*. *vulgaris* ssp. *maritima* could easily be distinguished, which was consistent with their morphology. Table beet and sugar beet accessions appeared to have clear genetic clusters with less admixture from other genetic clusters, which was also consistent with phenotypic observations. Interestingly, table beet accessions with complete genetic ancestry produce round-red roots. However, the table beet accessions with varying admixture from sugar beet ancestry form conical-red roots with banding patterns in the storage cambium tissues (Figure 1).

No distinct clusters of leaf and fodder beet ancestry were formed. Interestingly, leaf beets shared substantial genetic ancestry with the Mediterranean sea beet accessions (Figure 3B). The biennial-leaf beet accessions had more sugar beet ancestry than annual-leaf beet accessions. Fodder beet shared substantial genetic ancestry with table, sugar, and Mediterranean sea beet ancestry.

Accessions within the fodder beet cluster carrying different levels of sea beet ancestry may indicate varying degrees of breeding intensity. The fodder beet accessions with higher Mediterranean sea beet ancestry produce round beets. However, the accessions with higher sugar beet ancestry form sugar beet-like conical roots. Based on genetic ancestry, highly admixed sugar beet, table beet, and Mediterranean sea beet accessions were removed, and 245 accessions were kept for further analysis.

### Lineage-specific variation and phylogeny of the beet diversity panel

After removing the admixed accessions, genetic variation within each cultivated lineage and the wild sea beet populations was calculated. The number of SNPs was reduced to 10,052,949 and the INDELs to 1,260,345 across 245 samples. The Mediterranean sea beet accessions exhibited the highest number of SNPs and INDELs (9,751,509), followed by leaf beet (9,647,860), fodder beet (9,272,343), sugar beet (8,196,734), the Atlantic sea beet (7,456,809), and table beet accessions (7,425,689) (Supplementary Table 8).

The Mediterranean sea beet accessions shared the highest number of SNPs with leaf beet accessions (7,747,807 out of 9,520,600). In contrast, table beet accessions had the fewest SNPs in common with the Mediterranean sea beet accessions (5,879,955 out of 9,423,745). Interestingly, the table beet accessions also exhibited the lowest number of lineage-specific SNPs. Sugar beet accessions shared 6,258,918 SNPs with the Mediterranean sea beet accessions, but they also carried the highest number of lineage-specific SNPs (Supplementary Table 9).

The Atlantic sea beet sub-cluster shared the most SNPs with the Mediterranean sea beet accessions (5,845,172 out of 9,493,914). The fewest SNPs were shared with sugar beet (4,912,371) and table beet (4,364,533). Interestingly, all sub-clusters, except table beet, exhibited an abundance of lineage-specific SNPs. Additionally, table beet accessions shared the fewest common SNPs with the Atlantic sea beet (Supplementary Table 10).

Consistent with the PCA and admixture analysis results, the phylogenetic analysis revealed that cultivated beets are more closely related to Mediterranean sea beet than Atlantic sea beet. As expected, leaf beet accessions formed a sister clade with the Mediterranean sea beet accessions. All cultivated beets with storage roots formed one major clade. Within this ‘enlarged root’ clade, the table beet accessions formed a sister clade with fodder and sugar beet accessions. The fodder beet, conical-red table beet, and sugar beet accessions formed one clade. Commercial hybrids formed a separate clade, sharing the most common ancestor with the sugar beet germplasm resources from public breeding programs (Figure 3**Error****! Reference source not found.**C).

### Linkage disequilibrium and heterozygosity within the genetic sub-clusters of the beet diversity panel

We calculated linkage disequilibrium (LD) across the genome. LD decay between sequence variants averaged ∼2.5 kb. A rapid LD decay was more apparent in Mediterranean sea beet accessions (∼1.7 kb) than in cultivated beet lineages. Within the cultivated beet lineages, the table beet accessions had the largest LD blocks (∼5.9 kb) compared to sugar beet (∼4.6 kb), fodder beet (∼4.5 kb), and leaf beet accessions (∼3.0 kb) (Figure 4A, Table 2).

**Figure 4:**
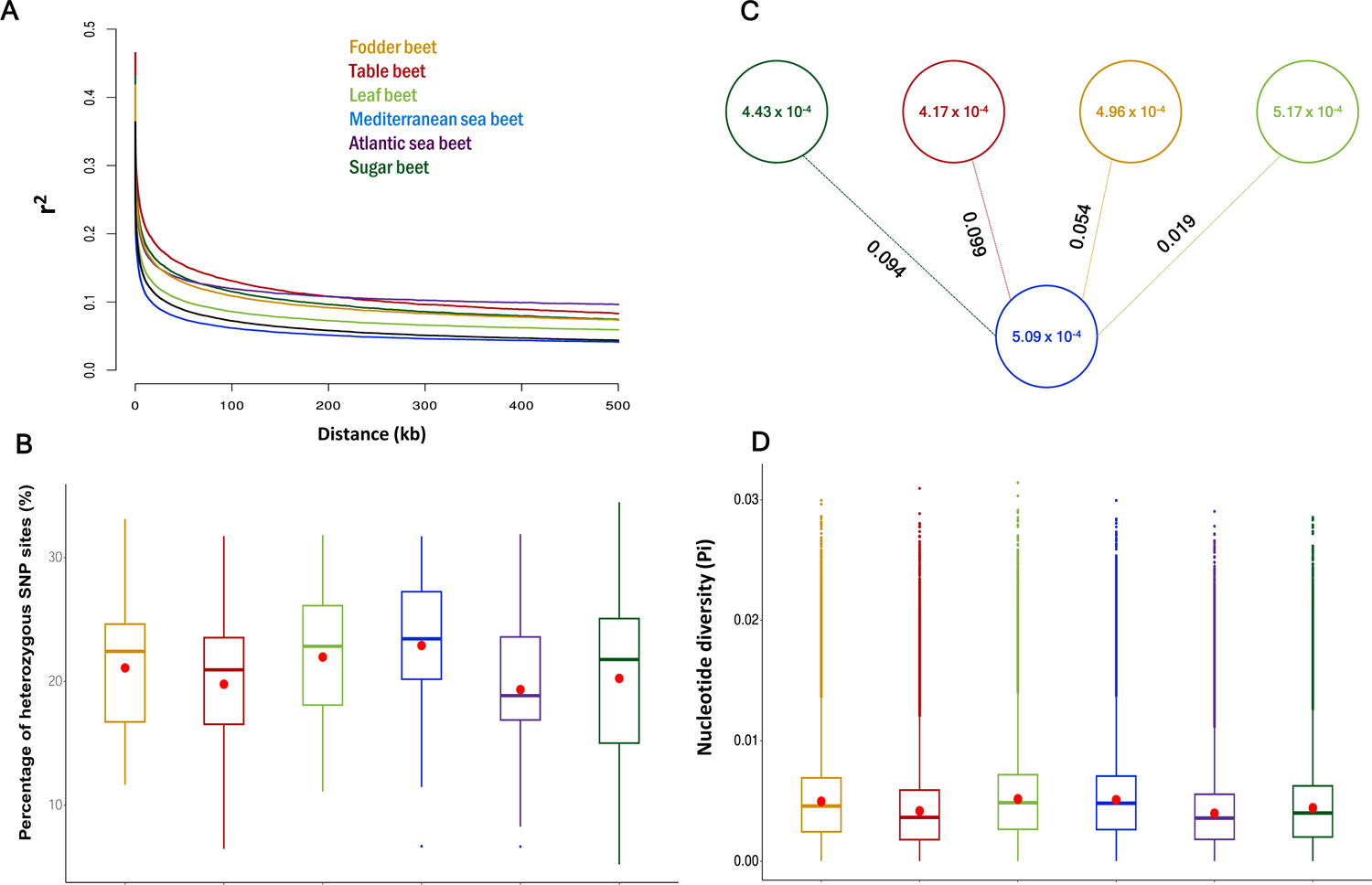
Linkage disequilibrium (LD) decay, percentage of heterozygous sites, population differentiation (FST), and nucleotide diversity within different genetic clusters of the Beta mini-core collection. The analyses were performed using whole genome SNP data of fodder beet (n=30), sugar beet (n=71), leaf beet (n=29), table beet (n=44), Atlantic (n=18), and Mediterranean sea beet (n=53) accessions. The color used in the analyses corresponds to the color codes used in PCA. (A) LD decay within each sub-cluster of the Beta mini-core collection. The Y-axis represents the pairwise correlation coefficient (r2) between two SNP markers and the X-axis represents the physical distance (kb) between corresponding SNP markers. (B) Overview of heterozygosity within each genetic cluster. Each box plot represents the percentage of heterozygous SNP sites within each genetic cluster. The red dot represents the mean value, while the centerline inside each box corresponds to the median. The lower and upper hinges of the box represent the 25th and 75th percentiles, respectively. Any data points located outside the whiskers are considered outliers. (C) Nucleotide diversity within both cultivated beet lineages and the Mediterranean sea beet accessions, along with pairwise FST values denoting the differentiation of cultivated beet lineages from the Mediterranean sea beet accessions. Each circle corresponds to the nucleotide diversity value of its respective group, and the value along each connecting line indicates the FST value between the two groups. (D) Overview of nucleotide diversity within each genetic cluster. Each box plot represents the nucleotide diversity within each genetic cluster. The red dot represents the mean value, while the centerline inside each box corresponds to the median. The lower and upper hinges of the box represent the 25th and 75th percentiles, respectively. Any data points located outside the whiskers are considered outliers.

**Table 2:** Linkage disequilibrium (LD) analysis across cultivated and wild *Beta* accessions.

| Type | Number of samples | Max. $r^2$ | Max. $r^2/2$ | Distance (kb) at Max. $r^2/2$ |
| --- | --- | --- | --- | --- |
| Fodder beet | 30 | 0.417 | 0.209 | 4.4 – 4.5 |
| Table beet | 44 | 0.465 | 0.232 | 5.8 – 5.9 |
| Leaf beet | 29 | 0.379 | 0.189 | 2.9 – 3.0 |
| Mediterranean sea beet | 53 | 0.343 | 0.171 | 1.6 – 1.7 |
| Atlantic sea beet | 18 | 0.418 | 0.209 | 3.7 – 3.8 |
| Sugar beet | 71 | 0.431 | 0.215 | 4.5 – 4.6 |
| Diversity Panel | 245 | 0.364 | 0.182 | 2.4 – 2.5 |

Among the cultivated lineages, table beet displayed large LD blocks across all chromosomes except chromosome 1 (Supplementary Figure 6). The largest LD blocks were on chromosome 2. In contrast, sugar beet accessions showed the largest LD blocks on chromosome 1 (Supplementary Figure 7). Leaf and fodder beet accessions displayed uniform LD decay across all nine chromosomes.

Cultivated and wild beets are out-crossing species due to an efficient self-incompatibility system. While commercial beet varieties are hybrids, inbred populations are commonly maintained by sib-mating. Therefore, we expected a considerable degree of heterozygosity. Across all the 290 accessions, 20.97% of the SNP sites were heterozygous (Supplementary Table 11). The highest average number of SNP variant sites were observed in the Mediterranean sea beet accessions (22.9%) followed by leaf beet (21.9%), fodder beet (21.1%), sugar beet (20.2%), table beet (19.8%) and Atlantic sea beet (19.3%) accessions (Supplementary Figure 8). In general, leaf beet and Mediterranean sea beet plants displayed higher heterozygosity rates than the rest of the mini-core collection (Supplementary Table 11). The sugar beet accessions showed the highest variation regarding heterozygosity rates, ranging from 5.24 % to 34.46 % (Figure 4B).

### Patterns of genomic variation between cultivated and wild beet accessions

We conducted pairwise FST calculations between cultivated and wild accessions to measure population differentiation. First, the Mediterranean sea beet accessions were the reference population due to their highest heterozygosity, suggesting their potential ancestral position for Atlantic and crop types. The highest mean fixation index was observed between the Atlantic and the Mediterranean sea beet accessions (*F*_ST_ = 0.103). Comparing Mediterranean sea beet and cultivated lineages revealed lower *F*_ST_ values (0.099 for table beet, 0.094 for sugar beet, 0.054 for fodder beet, and 0.019 for leaf beet) (Figure 4C, Table 3).

**Table 3:** Pairwise population differentiation (*F_ST_*) comparisons between the Mediterranean sea beet accessions and the rest of the *Beta* core collection.

|  | **Mediterranean sea beet** |  |  |
| --- | --- | --- | --- |
| **Populations** | **Mean $F_{ST}$** | **Weighted $F_{ST}$** | **Number of individuals** |
| Atlantic sea beet | 0.103 | 0.152 | 71 |
| Table beet | 0.099 | 0.137 | 97 |
| Sugar beet | 0.094 | 0.120 | 124 |
| Fodder beet | 0.054 | 0.072 | 83 |
| Leaf beet | 0.019 | 0.024 | 82 |

Then, we compared the Atlantic sea beet with the cultivated lineages, which resulted in higher *F*_ST_ values than the comparison with Mediterranean sea beet (Supplementary Table 12). The highest *F*_ST_ value was observed between table beet and Atlantic sea beet (0.197), followed by sugar beet (*F*_ST_ = 0.178), fodder beet (0.148), and leaf beet (0.116). The lowest *F*_ST_ values were found between the Mediterranean and the Atlantic sea beets.

We also assessed the nucleotide diversity within cultivated and wild beets (Supplementary Table 13). Surprisingly, the leaf beet accessions had the highest nucleotide diversity (π=5.17×10^-4^), followed by the Mediterranean sea beet (π=5.09×10^-4^), fodder beet (π=4.96×10^-4^), sugar beet (π=4.43×10^-4^) and table beet (π=4.17×10^-4^). The lowest nucleotide diversity was observed within the Atlantic sea beet accessions (π=3.96×10^-4^) (Figure 4C-D).

### Signatures in domesticated beet genomes pointing at genes under selection in a sucrose-storing root crop

The domestication and improvement process often leads to a purifying selection of genomic regions associated with favorable phenotypic traits. We expected that selection for highly heritable traits like sucrose content, bolting resistance, and root weight would leave selection footprints in the sugar beet genome. Therefore, we first employed XP-CLR and FST analyses, focusing on the top 5% of regions overlapping both approaches.

First, we compared sugar beet with the Mediterranean sea beet. We identified selective sweep regions in the EL10 genome spanning approximately 44.47 Mb (using XP-CLR) (Supplementary Figure 9) and 69.97 Mb (using *F*_ST_) (Supplementary Figure 10). In total, 30.27 Mb (5.32%) of the EL10 genomic region overlapped between both analyses, indicating putative selection footprints. The majority, approximately 54.16% (16.39 Mb), of the regions under selection was concentrated on chromosomes 1, 6 (Figure 5A-B), and 2, where also the longest and the highest number of sweeps were located (Table 4). Within these genome regions, we identified 1,741 potential domestication and improvement genes.

**Figure 5:**
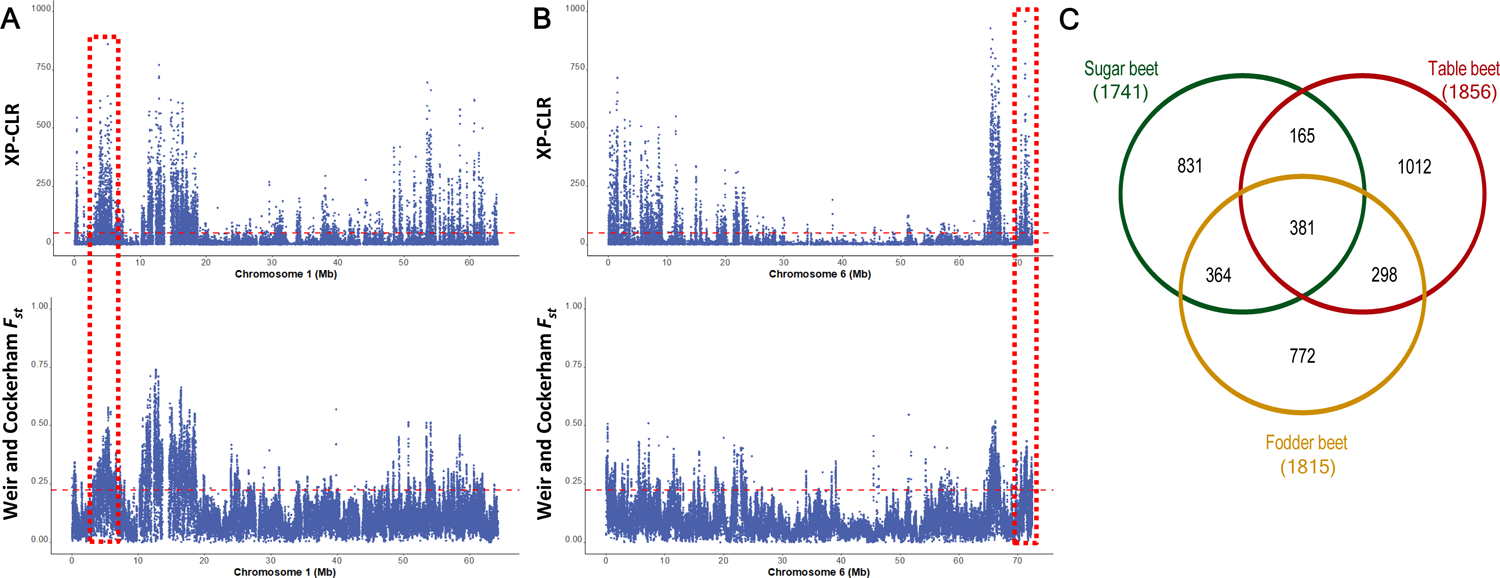
Selective sweeps in sugar beet and gene regions under selection in sugar, table, and fodder beets. (A) XP-CLR and FST across EL10 sugar beet chromosome 1 using 10 kb sliding window size and 1 kb step size while comparing sugar beet accessions with the Mediterranean sea beet accessions. The blue dots represent the XP-CLR and mean FST values in the upper and lower plots along the EL10 chromosome 1. The horizontal red line marks the empirical 95th percentile of XP-CLR and FST values. The red dashed box highlights the regions of interest carrying interesting candidate genes under strong selection. (B) XP-CLR and FST across EL10 sugar beet chromosome 6. (C) Venn diagram of shared and lineage-specific genes under selection in sugar beet, table beet, and fodder beet accessions across nine EL10 chromosomes.

**Table 4:** Summary statistics of selective sweep regions using XP-CLR and *F*_ST_ across EL10 chromosomes within the cultivated beet lineages forming a storage root. The overlapping genomic regions from XP-CLR and *F*_ST_ were considered the selective sweep regions. The table shows the total length (bp), maximum length (bp), average length (bp), and the number of selective sweep regions in sugar, table, and fodder beets across nine EL10 chromosomes.

|  | Sugar beet |  |  |  | Table beet |  |  |  | Fodder beet |  |  |  |
| --- | --- | --- | --- | --- | --- | --- | --- | --- | --- | --- | --- | --- |
| Chromosome | Total length (bp) | Maximum length (bp) | Average (bp) | Number of sweeps | Total length (bp) | Maximum length (bp) | Average (bp) | Number of sweeps | Total length (bp) | Maximum length (bp) | Average (bp) | Number of sweeps |
| 1 | 8,225,708 | 223,999 | 28,170 | 292 | 2,756,833 | 102,999 | 16,508 | 167 | 3,579,824 | 179,999 | 20,340 | 176 |
| 2 | 3,980,794 | 199,999 | 19,324 | 206 | 4,658,792 | 126,999 | 22,398 | 208 | 5,941,714 | 194,999 | 20,775 | 286 |
| 3 | 1,701,900 | 72,999 | 17,019 | 100 | 2,484,855 | 104,999 | 17,137 | 145 | 1,180,929 | 81,999 | 16,633 | 71 |
| 4 | 1,264,931 | 64,999 | 18,332 | 69 | 1,464,906 | 49,999 | 15,584 | 94 | 4,265,828 | 174,999 | 24,801 | 172 |
| 5 | 1,896,900 | 93,999 | 18,969 | 100 | 2,132,899 | 123,999 | 21,118 | 101 | 2,036,911 | 211,999 | 22,887 | 89 |
| 6 | 4,185,822 | 213,999 | 23,516 | 178 | 6,136,669 | 125,999 | 18,540 | 331 | 4,649,779 | 166,999 | 21,040 | 221 |
| 7 | 2,931,850 | 139,999 | 19,546 | 150 | 2,476,870 | 73,999 | 19,053 | 130 | 2,264,891 | 137,999 | 20,779 | 109 |
| 8 | 3,889,820 | 148,999 | 21,610 | 180 | 2,950,867 | 193,999 | 22,187 | 133 | 2,159,888 | 150,999 | 19,285 | 112 |
| 9 | 2,187,875 | 81,999 | 17,503 | 125 | 672,956 | 53,999 | 15,294 | 44 | 3,380,840 | 151,999 | 21,130 | 160 |
| Total | 30,265,600 |  |  | 1400 | 25,735,647 |  |  | 1353 | 29,460,604 |  |  | 1396 |
| Average | 3,362,844 |  | 20,443 | 156 | 2,859,516 |  | 18,647 | 150 | 3,273,400 |  | 20,852 | 155 |

Then, we compared table beet with Mediterranean sea beet. We identified selective sweep regions spanning approximately 45.53 Mb using *F*_ST_ and 65.68 Mb (XP-CLR) of the EL10 genome. In total, 25.74 Mb (4.53%) of the EL10 genomic region overlapped between both analyses, indicating putative selective sweep regions in table beet accessions. Nearly 53.41% (13.75 Mb) of the table beet genome under selection was localized on chromosomes 6, 2, and 8. Furthermore, the maximum length of selective sweeps and the number of sweeps were predominantly located on chromosomes 2, 6, and 8 (Table 4). Within these regions, we identified 1,856 potential candidate genes associated with selection or domestication in table beet accessions.

Next, we compared fodder beet accessions with the Mediterranean sea beet. Putative selective sweep regions spanned approximately 45.66 Mb and 72.77 Mb of the EL10 genome using *F*_ST_ and XP-CLR, respectively. In total, 29.46 Mb (5.18%) overlapped between both analyses. Nearly 50.43% (14.86 Mb) of the fodder beet genome under selection was situated on chromosomes 2, 6, and 4, where the maximum number of sweeps was also located (Table 4). Within these regions, we identified 1,815 genes. We reasoned that intensive selection for high root weight and against bolting during the past 200 years left its footprints in the cultivated root types (sugar beet, fodder beet, table beet).

Consequently, genes or genome regions involved in beet domestication and improvement traits should be conserved across these lineages. Among the genes within the candidate selection signature regions of these lineages, 381 genes were common (Figure 5C). As expected, sugar beet and fodder beet had the highest number of shared genes (745), followed by fodder beet/table beet (679), and sugar beet/table beet (546). Notably, table beet accessions exhibited the highest number of lineage-specific genes under selection (1,012), followed by sugar beet (831) and fodder beet (772).

Next, we searched for genes putatively involved in agronomic traits typical for a sucrose-storing biennial root crop (Supplementary Figure 11, Supplementary Figure 12). The huge number of DNA polymorphisms offered an opportunity to investigate nucleotide diversity between cultivated lineages and the Mediterranean sea beet on the gene level. First, we focussed on putative sucrose transporters. Sugar beet, table beet, and fodder beet accessions exhibited reduced nucleotide diversity within a homolog of the Arabidopsis *SUCROSE TRANSPORTER 4* (*SUT4)* on chromosome 1 when compared to leaf beets and Atlantic and Mediterranean sea beet (Figure 6A). Another sucrose transporter, *SUT2*, on chromosome 6 showed reduced nucleotide diversity exclusively in sugar beet accessions compared to all other sub-clusters (Figure 6B). However, no signatures of selective sweeps were detected in the sucrose transporter *BvTST2.1*, which is responsible for vacuolar sucrose uptake in sugar beet taproots ^38^, and *BvSUT1*, a member of the disaccharide transporter (DST) superfamily involved in the loading of sucrose into the phloem of sugar beet leaves ^39^.

**Figure 6:**
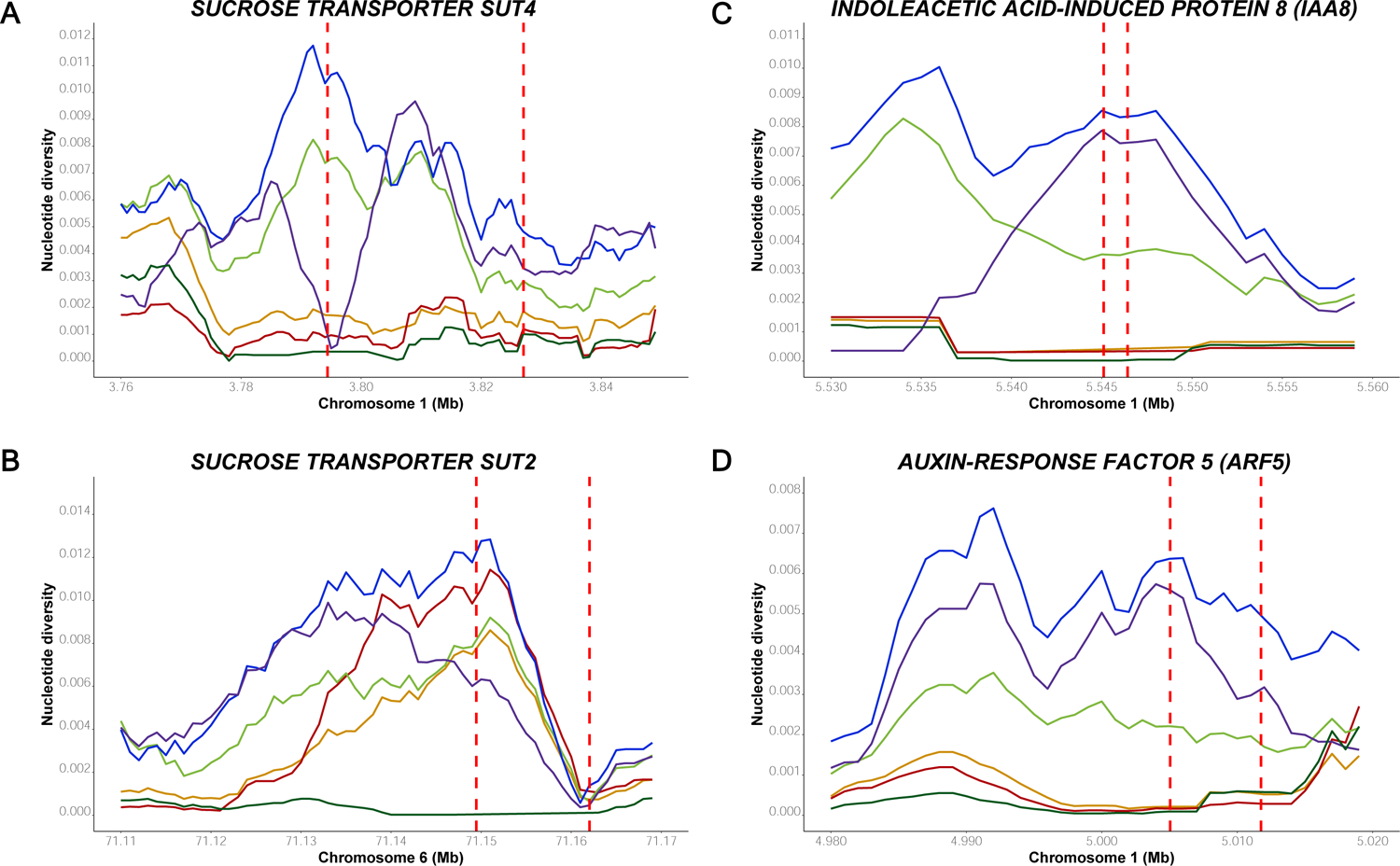
Nucleotide diversity within fodder beet, leaf beet, sugar beet, Atlantic sea beet, and Mediterranean sea beet accessions in and around the genomic regions housing genes involved in sucrose metabolism and auxin response. The color of the line corresponds to the color codes of the sub-clusters used in PCA. The Y-axis depicts the nucleotide diversity value, while the X-axis denotes the location along the chromosome. The vertical red lines represent the gene boundaries. (A) Nucleotide diversity values between different sub-clusters in the selective sweep region spanning the *SUT4* gene on chromosome 1. (B) Nucleotide diversity values between different sub-clusters in the selective sweep region spanning the *SUT2* gene on chromosome 6. (C) Nucleotide diversity values between different sub-clusters in the selective sweep region spanning *IAA8* gene on chromosome 1. (D) Nucleotide diversity values between different sub-clusters in the selective sweep region spanning *ARF5* gene on chromosome 1.

At the end of chromosome 1, homologs of two auxin-responsive genes from Arabidopsis are located within a region of reduced nucleotide diversity, *INDOLEACETIC ACID-INDUCED PROTEIN 8* (*IAA8*) (Figure 6C) and *AUXIN RESPONSE FACTOR 5* (*ARF5*) (Figure 6D). In Arabidopsis, it has been shown that auxin response factors play a crucial role in root development, lateral root formation, and the pattern formation of the root central vascular cylinder and vascular tissue. We hypothesize that cultivated beets were domesticated for their ability to form thickened roots. During the beet improvement process, breeders selectively increased the root size while simultaneously reducing root branching, also known as fangy roots—a typical characteristic of sea beet roots.

In cultivated beet accessions producing thickened storage roots, a complete loss of genetic diversity was observed within and around *IAA8* and *ARF5*, when compared to leaf beets, and Atlantic and Mediterranean sea beets. In Arabidopsis, it has been shown that *IAA8* is expressed in the developmental vasculature of the shoot apex, hypocotyl, and root tip, acting as a transcriptional repressor of auxin response and controlling lateral root formation ^40^. Interestingly, *ARF5*, widely known as *MONOPTEROS* (*MP*), encodes a transcription factor, *IAA24*, and plays a role in mediating embryo axis formation, vascular development, and the polar transport of auxins ^41, 42^.

Inositol transporters and raffinose synthase genes have been functionally characterized in beet. Therefore, we selected *INOSITOL TRANSPORTER 1* (*INT1*) ^20^ on chromosome 3 and *RAFFINOSE SYNTHASE 5* (*RS5*) ^43^ on chromosome 1. It is known that these genes are involved in post-harvest freezing tolerance, an important trait targeted during beet breeding. Raffinose, a trisaccharide, acts as a cryoprotectant in the pith of the storage taproot, and inositol is crucial for synthesizing raffinose. Interestingly, we found a reduced nucleotide diversity within *INT1* among sugar beet and table beet accessions compared to other sub-clusters (Supplementary Figure 11A). Additionally, reduced nucleotide diversity was observed within *RS5* only in sugar beets compared to all other sub-clusters (Supplementary Figure 11B).

Surprisingly, we found no evidence of selective sweeps within the ORFs of previously described flowering time genes such as *BvBTC1*, *BvFT1*, *BvFT2,* and *BvBBX19* ^44–48^. However, we did observe a reduced nucleotide diversity in the putative *FLC ( FLOWERING LOCUS C*) ortholog on chromosome 6 ^49^ and within a homolog of *SUPPRESSOR OF OVEREXPRESSION OF CO 1* (*SOC1*), an important floral integrator and a downstream target of *FLC* in Arabidopsis (Supplementary Figure 11C-D).

Table beets are known for their distinctive earthy taste in their roots, which is believed to have been intentionally selected for enhanced flavor. A QTL study in table beets revealed a strong association between high geosmin concentration and a locus on chromosome 8, with terpene synthesis genes located within this QTL region. Terpene synthase genes are known to be involved in geosmin biosynthesis (Hanson et al. 2021). Additionally, table beet accessions displayed a reduced nucleotide diversity within terpene synthase genes on chromosomes 7 and 8 (Supplementary Figure 12A-B).

## Discussion

We assembled a diversity panel of the genus *Beta* consisting of cultivated beet lineages and their wild progenitor *B. vulgaris* ssp. *maritima* (sea beet). This mini-core collection represents this species’ diverse geographical origin and phenotypic variation. Whole genome sequencing unraveled the genotypic diversity within and between cultivated and wild lineages, the gene flow between subspecies, and the footprints of selection during domestication and breeding. Within the genomes of cultivated beets, putative selective sweeps of extremely low genetic diversity were identified. These regions harbor candidate genes with a putative role in storage root formation and sucrose storage.

We applied PCA and admixture analyses to analyse the relatedness between sea beet accessions. Consistent with prior studies, sea beets formed two distinct clusters (Mediterranean and Atlantic) ^19, 20, 50^. This divergence is likely a result of the geographical separation between these regions, leading to limited genetic exchange or gene flow among the wild accessions. Consistent with other studies, the Atlantic sea beets, especially those from Ireland, were most distantly related to the sugar beet line EL10, the donor of the reference genome sequence ^20^. PCA and phylogenetic analyses revealed remarkable proximity between the Mediterranean sea beet and cultivated beets, suggesting that Mediterranean sea beet is the immediate ancestor of cultivated beets. Moreover, a combination of PCA, admixture, shared SNPs, and *F_st_* data revealed for the first time that leaf beets, among the cultivated lineages, are genetically closest to the Mediterranean sea beets. Remarkably, this finding aligns with historical data, indicating that ancient Greek and Roman civilizations used the leaves of vegetable beet in their culinary practices and medicinal remedies ^1, 5^. Interestingly, all cultivated beets with enlarged thickened roots lie in one clade, which suggests that the beets producing thickened roots share a more recent common ancestor different from leaf beets. Noteworthy, all modern hybrids cluster closely in one subclade within the sugar beet clade, consistent with previous findings ^19^.

Then, we analysed the relatedness among cultivated lineages. Sugar and table beet accessions exhibited distinct genetic clusters, with only a few accessions with genetic admixtures from other ancestries. This result reflects strong selection, intense breeding for diverse breeding aims, and genetic drift due to reproductive isolation within these cultivated lineages. The divergence between these two lineages corresponds to previous studies ^8, 51, 52^. The effective population size further indicated these patterns, and a recent rapid bottleneck was identified in the past 500 years within both sugar and table beet cultivated lineages (Supplementary Figure 13). Documentary evidence also supports this finding, indicating that beet roots were not used as livestock feed or as a vegetable before the 16^th^ century ^5, 53^.

A small subset of sugar beet accessions exhibited highly admixed ancestry, primarily from Mediterranean sea beet, corresponding to their diverging phenotypic characteristics like root fanginess. These accessions originated from Turkey and Greece, which are recognized as the primary centers of beet domestication ^5, 8, 19^. This could mean that beet from these regions were first domesticated from the wild Mediterranean sea beet, and later, they have not undergone further admixture or intense selection. Alternatively, this finding could point to inaccurate passport data.

Most table beets originating from Greece, Soviet Union, and Turkey displayed a distinct, clean ancestry, which indicates extensive selection and breeding efforts. The conical roots and an alternating red and white pattern in the parenchyma and vascular tissue, observed in certain table beet accessions due to betalain accumulation, could be attributed to the introgression of genomic regions governing the conical root shape and sucrose content from sugar beet ancestry, as revealed by admixture analysis.

Based on our PCA analysis, fodder beet, and leaf beets were considered intermediate between table and sugar beets, which was also supported by the admixture analysis, where no distinct genetic clusters were found for these cultivated lineages. Notably, among fodder beets, those accessions with a higher percentage of sugar beet ancestry, characterized by their sugar beet-like conical roots, originate from Europe. It is widely acknowledged that the ‘White Silesian’ sugar beet progenitor was derived from sucrose-storing ‘Runkelrüben’ lineages. Thus, our findings align with the existing knowledge regarding the genetic relationships between fodder and sugar beets ^2, 8, 54^. Furthermore, the fodder beet accessions with round roots originating from Syria, Iran, Turkey, and China exhibited a notably higher percentage of Mediterranean sea and table beet ancestry than sugar beet (**Error! Reference source not found.**B). This observation leads to the tempting speculation that fodder beets with both round and flat roots emerged in the Mediterranean region, and the development of such fodder beets could be attributed to the spontaneous or deliberate hybridization events between table beets and Mediterranean sea or leaf beet accessions.

Cultivated beets forming a storage root displayed reduced genetic diversity compared to Mediterranean sea beet, likely due to the bottleneck effect resulting from domestication and artificial selection. The fixation index (*F_st_*), a measure of population differentiation, ranged from 0.019 to 0.099 in leaf beet and table beet accessions. This range is comparable to the *F_st_* difference between *Spinacia oleracea* and *S*. *turkestanica* (0.030)^55^ but considerably lower than the difference observed between lettuce species (0.305)^56^ and cultivated and wild rice (0.30)^57^. The lower *F_st_* values could be attributed to the high number of shared SNPs between leaf and fodder beets and Mediterranean sea beets.

We examined the impact of domestication and artificial selection at the gene level, identifying genome regions with reduced genetic diversity in cultivated beets. According to Fst and XP-CLR analyses, shared regions were considered areas under strong selection. Domestication and intensive breeding likely contributed to the decreased genetic diversity in these regions, with the implicated genes influencing these processes. We found 30.27 Mb of genome region with very low diversity or complete fixation in sugar beet but with high diversity in the Mediterranean sea beet accessions. They can be interpreted as selective sweeps due to strong selection over the past 200 years. Notably, all agronomically important characters can be selected in root-type forms before flowering, which may explain the high success rate of mass selection.

In the beet lineages characterized by enlarged taproots, 381 shared genes were found to be under selection. Homologs of two auxin-responsive genes from Arabidopsis (*IAA8, ARF5*) are potential candidate genes involved in the formation of storage roots. Auxin plays a pivotal regulatory role in diverse developmental processes, encompassing root and stem growth, vascular differentiation, embryo patterning, and lateral branching patterns in the root ^58^. In Arabidopsis, *IAA8*, encoding a transcriptional repressor of the auxin response, interacts with the TIR1 auxin receptor and ARF (<u>a</u>uxin response factors) transcription factors in the nucleus, thereby governing lateral root formation. Transgenic lines overexpressing *IAA8* exhibited a significantly lower number of lateral roots than the wild type, whereas loss-of-function mutant lines of *IAA8* displayed a significantly higher number of lateral roots ^40^. A study identified potential interactions of the IAA8 protein with ten ARFs, including ARF5 ^59^.

We hypothesize that the interaction between Aux/IAA transcriptional repressors and auxin response factor genes is instrumental in reducing lateral branching and increasing root size. In beets, it has been demonstrated that the transition from primary growth to secondary growth involves transient changes in the levels of auxin (IAA), active cytokinins (CKs), and abscisic acid (ABA). This suggested their involvement in regulating the initiation of secondary growth from the cambial rings^60^.

In addition to these genes, we observed reduced genetic diversity in a *WUSCHEL-RELATED HOMEOBOX 4* (*WOX4*) gene on chromosome 1, which is under artificial selection in sugar, table, and fodder beets. In Arabidopsis, cambium cell-specific transcript profiling demonstrated that *WOX4* is one of the two key master regulators of cambium activity. It plays a role in xylem cell expansion and stem cell regulation and promotes cambial cell proliferation in Arabidopsis roots ^61^.

We were also interested in genes regulating sucrose storage. We identified two sucrose transporter genes on chromosomes 1 and 6. The consistent pattern of significantly reduced genetic diversity in both genes suggests their potential involvement in high sucrose storage in sugar beet roots.

Our study sheds light on the genetic diversity and composition of cultivated and wild beets, revealing the impact of domestication and intensive breeding on the beet genome. Future research will provide deeper insights, including haplotype analyses, and building and functional analyses of candidate genes. The *Beta* mini-core collection and the genome sequences are important resources that will allow genome-wide association studies and allele mining to introduce new allelic variation to the gene pool of cultivated beets.

## Acknowledgments

We thank Paul Galewski (Department of Plant, Soil, and Microbial Science, Michigan State University), for providing the EL10.2_2 genome assembly. We thank Brigitte Neidhardt-Olf, Bettina Rohardt, Ines Schütt, and Monika Bruisch for their technical assistance. We gratefully acknowledge financial support from the German Science Foundation (Deutsche Forschungsgemeinschaft, DFG) – Project Number 400993799 (Project 2 within the Research Training Group 2501 Translational Evolutionary Research, https://gepris.dfg.de/gepris/projekt/400993799).

## Supplementary tables

**Supplementary Table 1:** Sea beet and cultivated beet accessions used in this study.

**Supplementary Table 2:** Summary statistics of the paired-end Illumina raw reads without trimming from 290 accessions.

**Supplementary Table 3:** Summary statistics of the paired-end Illumina trimmed and cleaned reads from 290 accessions.

**Supplementary Table 4:** Summary statistics of the sample information, total mapped reads (%), sequencing depth and genome coverage (%) across 290 accessions.

**Supplementary Table 5:** Summary statistics of sequence variants (SNPs and INDELs) in 290 beet accessions compared to the EL10 reference genome.

**Supplementary Table 6:** Summary statistics of the number of SNPs and INDELs with their predicted effect using VEP across 290 accessions.

**Supplementary Table 7:** Summary statistics of the number of SNPs and INDELs within coding sequences with their predicted effect using VEP across 290 accessions.

**Supplementary Table 8:** Summary statistics of sequence variants (SNPs and INDELs) within each sub-cluster compared to the EL10 reference genome.

**Supplementary Table 9:** Pairwise shared variants between the Mediterranean sea beet accessions and the other sub-clusters.

**Supplementary Table 10:** Pairwise shared variants between the Atlantic sea beet accessions and the other sub-clusters.

**Supplementary Table 11:** Summary statistics of the percentage of heterozygous SNPs in each accession.

**Supplementary Table 12:** Summary statistics of pairwise comparisons of population differentiation (Fst) between the Atlantic sea beet accessions and the other sub-clusters.

**Supplementary Table 13:** Summary statistics of nucleotide diversity within each genetic sub-cluster.

## Supplementary figures

**Supplementary Figure 1:** Overview of the bioinformatics workflow used in the study. The Illumina short-reads were mapped to the EL10.2_2 genome assembly using BWA-MEM.

**Supplementary Figure 2:** A bar plot illustrating the number of variants, including SNPs and INDELs, across all nine EL10 chromosomes in the Beta mini-core collection of 290 accessions. The Y-axis represents the total number of variants, and the X-axis corresponds to the nine EL10 chromosomes. Chromosome 9 exhibits the minimum number of variants, approximately 1.1 million, while Chromosome 5 displays the maximum number, around 1.5 million variants. On average, each chromosome harbors approximately 1.3 million variants.

**Supplementary Figure 3:** Principal component analysis (PCA) of 290 beet accessions comparing second and third principal components. PC2 and PC3 represent the second and third components, accounting for 3.08% and 2.50% of the total variation, respectively.

**Supplementary Figure 4:** Principal component analysis (PCA) of 199 cultivated beet accessions, including sugar, table, fodder, and leaf beet accessions. PC1 and PC2 denote the first and second components, explaining 4.39% and 3.25% of the total variation, respectively.

**Supplementary Figure 5:** Cross-validation errors for each K in 10 replicates tested using model-based clustering in ADMIXTURE, incorporating varying numbers of ancestral kinships (K=1 and K=10). The Y-axis represents the cross-validation error value, while the X-axis corresponds to the Kth ancestry. The plot represents the cross-validation errors from 10 replicates at each K. Notably, the minimum cross-validation error and the variation within 10 replicates were observed at K=4.

**Supplementary Figure 6:** Linkage disequilibrium (LD) decay among distinct genetic clusters of the Beta mini-core collection across all nine chromosomes. Accessions with low admixture are depicted in black. LD is plotted across nine chromosomes (1 to 9). Within each plot, the Y-axis represents the pairwise correlation coefficient (r2) between two SNP markers, and the X-axis represents the physical distance (kb) between corresponding SNP markers.

**Supplementary Figure 7:** Linkage disequilibrium (LD) decay comparisons among nine chromosomes within each Beta mini-core collection genetic cluster. LD decay comparisons are depicted within sugar beet, table beet, fodder beet, leaf beet, Mediterranean ssp. *maritima*, Atlantic sea beet, and 245 accessions with low admixture from other genetic ancestries. LD is collectively plotted across all nine chromosomes (1 to 9) within each distinct genetic cluster. In each plot, the Y-axis represents the pairwise correlation coefficient (r2) between two SNP markers, and the X-axis represents the physical distance (kb) between corresponding SNP markers. The colors marked in the legend represent nine chromosomes.

**Supplementary Figure 8:** The frequency distribution plots of the percentage of heterozygous SNPs within distinct genetic clusters. The frequency distribution of heterozygous SNPs within each genetic cluster, such as Mediterranean sea beet, leaf beet, fodder beet, sugar beet, table beet, and Atlantic sea beet, is plotted. Six classes were calculated, ranging from 5-10%, 10-15%, 15-20%, 20-25%, 25-30% and 30-35%. In each plot, the Y-axis represents the frequency of individual accessions belonging to the representative classes, plotted on the X-axis. The colors distinguish between the six different classes.

**Supplementary Figure 9:** XP-CLR revealed selection sweeps in sugar beets across all nine EL10 chromosomes. XP-CLR across the nine EL10 chromosomes using 10 kb sliding window size and 1 kb step size while comparing sugar beet accessions with the Mediterranean sea beet accessions. The blue dots indicate XP-CLR values along the nine EL10 chromosomes, while the horizontal red line denotes the empirical 95th percentile threshold. According to XP-CLR, genomic regions above this threshold are considered putative regions under selection. The Y-axis represents the XP-CLR values ranging from 0 to 1000, and the X-axis illustrates the chromosome length in Mb.

**Supplementary Figure 10:** Population differentiation using fixation index (FST) between sugar beet and Mediterranean sea beet accessions across all nine EL10 chromosomes. The FST calculations employed 10 kb sliding window size and 1 kb step size while comparing sugar beet accessions with the Mediterranean sea beet accessions. The blue dots on the graphs indicate FST values along the nine EL10 chromosomes, while the horizontal red line represents the empirical 95th percentile threshold. Genomic regions above this threshold are considered highly differentiated regions between sugar beet and Mediterranean sea beet, as FST indicates. The Y-axis represents FST values ranging from 0 to 1, where 0 signifies no genetic differentiation and 1 indicates complete genetic differentiation between the two populations. The X-axis illustrates the chromosome length in Mb.

**Supplementary Figure 11:** Nucleotide diversity values within various genetic clusters, including leaf and fodder beets, in the regions of interest encompassing selective sweeps, with a specific focus on genes involved in cold tolerance during post-harvest storage and vernalization pathway in flowering time control. The colors represent different genetic clusters based on beet type, representing the wild accession from the Atlantic (purple) and the Mediterranean (blue), sugar beet (dark green), table beet (red), fodder beet (orange), and leaf beet (light green) accessions. The Y-axis illustrates the nucleotide diversity value, while the X-axis denotes the location along the chromosome. The vertical red lines represent the gene boundaries of the predicted gene models. The nucleotide diversity values across different clusters are graphically represented for (A) inositol transporter 1 on chromosome 2, (B) raffinose synthase 5 on chromosome 1, (C) FLOWERING LOCUS C (FLC) on chromosome 6, and (D) SUPPRESSOR OF OVEREXPRESSION OF CO 1 (SOC1) on chromosome 1.

**Supplementary Figure 12:** Nucleotide diversity values within various genetic clusters, including leaf and fodder beets, in the regions of interest encompassing selective sweeps, with a specific focus on genes associated with earthy flavor and plant immunity. The colors represent different genetic clusters based on beet type, representing the wild accession from the Atlantic (purple) and the Mediterranean (blue), sugar beet (dark green), table beet (red), fodder beet (orange), and leaf beet (light green) accessions. The Y-axis illustrates the nucleotide diversity value, while the X-axis denotes the location along the chromosome. The vertical red lines represent the gene boundaries of the predicted gene models. The nucleotide diversity values across different clusters are graphically represented for (A) terpene synthase on chromosome 7, (B) terpene synthase on chromosome 8, (C) a protein kinase involved in rust resistance in Arabidopsis on chromosome 2, and (D) three consecutive genes associated with plant immunity on chromosome 6.

**Supplementary Figure 13:** Estimated historical effective population size (Ne) of genetic clusters in the Beta mini-core collection using the program MSMC2. The X-axis represents the years on a logarithmic scale, while the Y-axis represents the effective population size. The mutation rate of 1.25e-8 was used. The colors represent different genetic clusters according to their beet type: wild accession from the Atlantic (purple), wild accession from the Mediterranean (blue), sugar beet (dark green), table beet (red), fodder beet (orange), and leaf beet (light green) accessions.

## References

1. Biancardi, E., Panella, L. W. & McGrath, J. M. Beta maritima: The Origin of Beets. Cham: Springer International Publishing 2, XXVI, 284, 10.1007/978-3-030-28748-1 (2020).

2 Scott, R. & Cooke, D. The sugar beet crop. (Chapman & Hall, 1993).

3 Hatlestad, G. J. et al. The beet *R* locus encodes a new cytochrome P450 required for red betalain production. Nature Genetics 44, 816–820, doi:10.1038/ng.2297 (2012).

4 Hatlestad, G. J. et al. The beet *Y* locus encodes an anthocyanin MYB-like protein that activates the betalain red pigment pathway. Nature Genetics 47, 92–96, doi:10.1038/ng.3163 (2015).

5 Goldman, I. L. & Janick, J. Evolution of Root Morphology in Table Beet: Historical and Iconographic. Front Plant Sci 12, 689926, doi:10.3389/fpls.2021.689926 (2021).

6 Goldman, I. L. & Austin, D. Linkage among the R, Y and BI loci in table beet. Theoretical and Applied Genetics 100, 337–343, doi:10.1007/s001220050044 (2000).

7 Gayon, J. & Zallen, D. T. The Role of the Vilmorin Company in the Promotion and Diffusion of the Experimental Science of Heredity in France, 1840–1920. Journal of the History of Biology 31, 241–262, doi:10.1023/A:1004335619901 (1998).

8 Galewski, P. & McGrath, J. M. Genetic diversity among cultivated beets (*Beta vulgaris*) assessed via population-based whole genome sequences. BMC Genomics 21, 189, doi:10.1186/s12864-020-6451-1 (2020).

9 Melzer, S., Müller, A. E. & Jung, C. in Genomics of Plant Genetic Resources: Volume 2. Crop productivity, food security and nutritional quality (eds Roberto Tuberosa, Andreas Graner, & Emile Frison) 3–26 (Springer Netherlands, 2014).

10 Melzer, S., Müller, A. E. & Jung, C. in *Genomics of Plant Genetic Resources: Volume 2*. Crop productivity, food security and nutritional quality 3–26 (Springer, 2013).

11 Artschwager, E. Anatomy of the vegetative organs of the sugar beet. J. agric. Res 33, 143–176 (1926).

12. Frese, L., Desprez, B. & Ziegler, D. Potential of genetic resources and breeding strategies for base-broadening in Beta. (2000).

13 Arumuganathan, K. & Earle, E. D. Estimation of nuclear DNA content of plants by flow cytometry. Plant Molecular Biology Reporter 9, 415–415, doi:10.1007/BF02672017 (1991).

14 Flavell, R. B., Bennett, M. D., Smith, J. B. & Smith, D. B. Genome size and the proportion of repeated nucleotide sequence DNA in plants. Biochemical Genetics 12, 257–269, doi:10.1007/BF00485947 (1974).

15 Dohm, J. C. et al. The genome of the recently domesticated crop plant sugar beet (Beta vulgaris). Nature 505, 546–549, doi:10.1038/nature12817 http://www.nature.com/nature/journal/v505/n7484/abs/nature12817.html#supplementary-information (2014).

16 McGrath, J. M. et al. A contiguous de novo genome assembly of sugar beet EL10 (*Beta vulgaris* L.). DNA Research 30, dsac033, doi:10.1093/dnares/dsac033 (2023).

17 Capistrano-Gossmann, G. G. et al. Crop wild relative populations of *Beta vulgaris* allow direct mapping of agronomically important genes. Nature Communications 8, 15708, doi:10.1038/ncomms15708 (2017).

18 Rodríguez Del Río, Á., et al. Genomes of the wild beets *Beta patula* and *Beta vulgaris* ssp. *maritima*. The Plant journal: for cell and molecular biology 99, 1242–1253, doi:10.1111/tpj.14413 (2019).

19 Sandell, F. L. et al. Genomic distances reveal relationships of wild and cultivated beets. Nature Communications 13, 2021, doi:10.1038/s41467-022-29676-9 (2022).

20 Felkel, S., Dohm, J. C. & Himmelbauer, H. Genomic variation in the genus *Beta* based on 656 sequenced beet genomes. Scientific Reports 13, 8654, doi:10.1038/s41598-023-35691-7 (2023).

21. Andrews, S. FastQC: a quality control tool for high throughput sequence data. Cambridge, UK: Babraham Bioinformatics, Babraham Institute (2010).

22 Ewels, P., Magnusson, M., Lundin, S. & Käller, M. MultiQC: summarize analysis results for multiple tools and samples in a single report. Bioinformatics 32, 3047–3048, doi:10.1093/bioinformatics/btw354 (2016).

23 Bolger, A. M., Lohse, M. & Usadel, B. Trimmomatic: a flexible trimmer for Illumina sequence data. Bioinformatics 30, 2114–2120, doi:10.1093/bioinformatics/btu170 (2014).

24 Li, H. & Durbin, R. Fast and accurate long-read alignment with Burrows–Wheeler transform. Bioinformatics 26, 589–595, doi:10.1093/bioinformatics/btp698 (2010).

25 Li, H. et al. The Sequence Alignment/Map format and SAMtools. Bioinformatics 25, 2078–2079, doi:10.1093/bioinformatics/btp352 (2009).

26 Van der Auwera, G. A. et al. From FastQ data to high confidence variant calls: the Genome Analysis Toolkit best practices pipeline. Curr Protoc Bioinformatics 43, 11 10 11-11 10 33, doi:10.1002/0471250953.bi1110s43 (2013).

27 McKenna, A. et al. The Genome Analysis Toolkit: a MapReduce framework for analyzing next-generation DNA sequencing data. Genome Res 20, 1297–1303, doi:10.1101/gr.107524.110 (2010).

28 McLaren, W. et al. The Ensembl Variant Effect Predictor. Genome Biology 17, 122, doi:10.1186/s13059-016-0974-4 (2016).

29 Krzywinski, M. et al. Circos: an information aesthetic for comparative genomics. Genome Res 19, 1639–1645, doi:10.1101/gr.092759.109 (2009).

30 Purcell, S. et al. PLINK: a tool set for whole-genome association and population-based linkage analyses. Am J Hum Genet 81, 559–575, doi:10.1086/519795 (2007).

31 Zheng, X. et al. A high-performance computing toolset for relatedness and principal component analysis of SNP data. Bioinformatics 28, 3326–3328, doi:10.1093/bioinformatics/bts606 (2012).

32 Alexander, D. H. & Lange, K. Enhancements to the ADMIXTURE algorithm for individual ancestry estimation. BMC Bioinformatics 12, 246, doi:10.1186/1471-2105-12-246 (2011).

33 Behr, A. A., Liu, K. Z., Liu-Fang, G., Nakka, P. & Ramachandran, S. pong: fast analysis and visualization of latent clusters in population genetic data. Bioinformatics 32, 2817–2823, doi:10.1093/bioinformatics/btw327 (2016).

34 Zhang, C., Dong, S.-S., Xu, J.-Y., He, W.-M. & Yang, T.-L. PopLDdecay: a fast and effective tool for linkage disequilibrium decay analysis based on variant call format files. Bioinformatics 35, 1786–1788 (2019).

35 Danecek, P. et al. The variant call format and VCFtools. Bioinformatics 27, 2156–2158 (2011).

36 Chen, H., Patterson, N. & Reich, D. Population differentiation as a test for selective sweeps. Genome Res 20, 393–402, doi:10.1101/gr.100545.109 (2010).

37 Schiffels, S. & Wang, K. MSMC and MSMC2: The Multiple Sequentially Markovian Coalescent. Methods Mol Biol 2090, 147–166, doi:10.1007/978-1-0716-0199-0_7 (2020).

38 Jung, B. et al. Identification of the transporter responsible for sucrose accumulation in sugar beet taproots. Nature Plants 1, 14001, doi:10.1038/nplants.2014.1 (2015).

39 Nieberl, P. et al. Functional characterisation and cell specificity of *BvSUT1*, the transporter that loads sucrose into the phloem of sugar beet (*Beta vulgaris* L.) source leaves. Plant Biology 19, 315–326, 10.1111/plb.12546 (2017).

40 Arase, F. et al. IAA8 Involved in Lateral Root Formation Interacts with the TIR1 Auxin Receptor and ARF Transcription Factors in *Arabidopsis*. PLOS ONE 7, e43414, doi:10.1371/journal.pone.0043414 (2012).

41 Berleth, T. & Jürgens, G. The role of the monopteros gene in organising the basal body region of the Arabidopsis embryo. Development 118, 575–587, doi:10.1242/dev.118.2.575 (1993).

42 Przemeck, G. K. H., Mattsson, J., Hardtke, C. S., Sung, Z. R. & Berleth, T. Studies on the role of the Arabidopsis gene *MONOPTEROS* in vascular development and plant cell axialization. Planta 200, 229–237, doi:10.1007/BF00208313 (1996).

43 Keller, I. et al. Cold-Triggered Induction of ROS- and Raffinose Metabolism in Freezing-Sensitive Taproot Tissue of Sugar Beet. Front Plant Sci 12, 715767, doi:10.3389/fpls.2021.715767 (2021).

44 Dally, N., Eckel, M., Batschauer, A., Höft, N. & Jung, C. Two CONSTANS-LIKE genes jointly control flowering time in beet. Scientific Reports 8, 16120, doi:10.1038/s41598-018-34328-4 (2018).

45 Höft, N., Dally, N., Hasler, M. & Jung, C. Haplotype variation of flowering time genes of sugar beet and its wild relatives and the impact on life cycle regimes. Frontiers in Plant Science 8, 2211 (2018).

46 Pin, P. A. et al. An Antagonistic Pair of FT Homologs Mediates the Control of Flowering Time in Sugar Beet. Science 330, 1397–1400, doi:10.1126/science.1197004 (2010).

47 Pin, Pierre A. et al. The Role of a Pseudo-Response Regulator Gene in Life Cycle Adaptation and Domestication of Beet. Current Biology 22, 1095–1101, 10.1016/j.cub.2012.04.007 (2012).

48 Dally, N., Xiao, K., Holtgräwe, D. & Jung, C. The *B2* flowering time locus of beet encodes a zinc finger transcription factor. Proceedings of the National Academy of Sciences 111, 10365–10370, doi:10.1073/pnas.1404829111 (2014).

49 Reeves, P. A. et al. Evolutionary conservation of the FLOWERING LOCUS C-mediated vernalization response: Evidence from the sugar beet (Beta vulgaris). Genetics 176, 295–307, doi:10.1534/genetics.106.069336 (2007).

50 Richards, C. M., Reeves, P. A., Fenwick, A. L. & Panella, L. Genetic structure and gene flow in *Beta vulgaris* subspecies *maritima* along the Atlantic coast of France. Genetic Resources and Crop Evolution 61, 651–662, doi:10.1007/s10722-013-0066-1 (2014).

51 Mangin, B. et al. Breeding patterns and cultivated beets origins by genetic diversity and linkage disequilibrium analyses. Theoretical and Applied Genetics 128, 2255–2271, doi:10.1007/s00122-015-2582-1 (2015).

52 Andrello, M., Henry, K., Devaux, P., Desprez, B. & Manel, S. Taxonomic, spatial and adaptive genetic variation of *Beta* section *Beta*. Theoretical and Applied Genetics 129, 257–271 (2016).

53 Nottingham, S. Beetroot. Self published (2004).

54 Fischer, H. E. Origin of the ‘Weisse Schlesische Rübe’(white Silesian beet) and resynthesis of sugar beet. Euphytica 41, 75–80 (1989).

55 Cai, X. et al. Genomic analyses provide insights into spinach domestication and the genetic basis of agronomic traits. Nature Communications 12, 7246, doi:10.1038/s41467-021-27432-z (2021).

56 Wei, T. et al. Whole-genome resequencing of 445 Lactuca accessions reveals the domestication history of cultivated lettuce. Nature Genetics 53, 752–760, doi:10.1038/s41588-021-00831-0 (2021).

57 Huang, X. et al. A map of rice genome variation reveals the origin of cultivated rice. Nature 490, 497–501 (2012).

58 Du, Y. & Scheres, B. Lateral root formation and the multiple roles of auxin. Journal of experimental botany 69, 155–167 (2018).

59 Wang, J., Yan, D.-W., Yuan, T.-T., Gao, X. & Lu, Y.-T. A gain-of-function mutation in *IAA8* alters Arabidopsis floral organ development by change of jasmonic acid level. Plant Molecular Biology 82, 71–83, doi:10.1007/s11103-013-0039-y (2013).

60 Jammer, A. et al. Early-stage sugar beet taproot development is characterized by three distinct physiological phases. Plant Direct 4, e00221, 10.1002/pld3.221 (2020).

61 Zhang, J. et al. Transcriptional regulatory framework for vascular cambium development in Arabidopsis roots. Nature Plants 5, 1033–1042, doi:10.1038/s41477-019-0522-9 (2019).

