## Supplementary Figures for "Signatures in domesticated beet genomes pointing at genes under selection in a sucrose-storing root crop"

### Slide 1
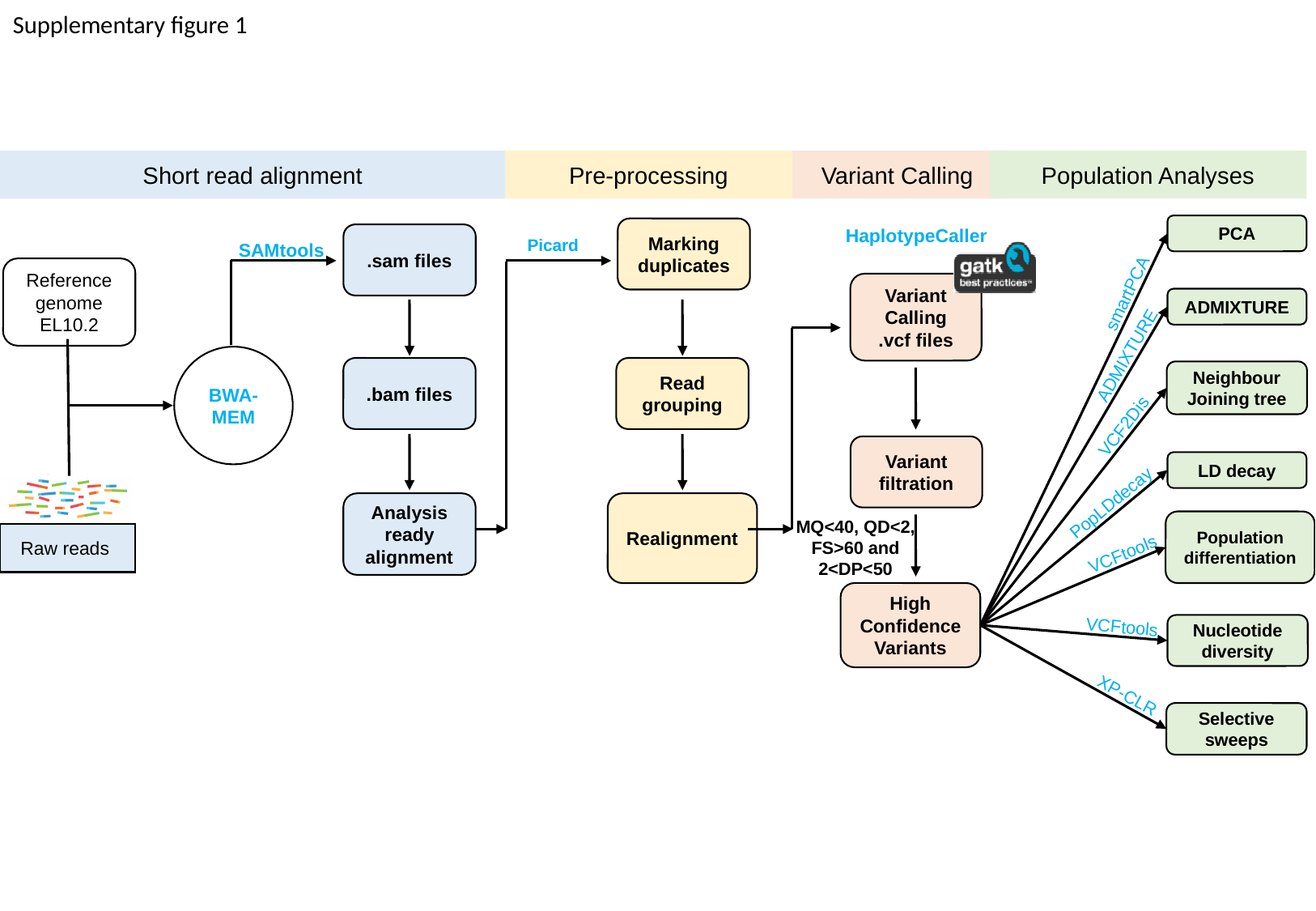

Supplementary figure 1
Variant Calling
Short read alignment
Pre-processing
HaplotypeCaller
Marking duplicates
SAMtools
.sam files
Picard
Reference genome EL10.2
Variant Calling
.vcf files
BWA-MEM
.bam files
Read grouping
Analysis ready alignment
Realignment
Raw reads
High Confidence Variants
Population Analyses
PCA
smartPCA
ADMIXTURE
ADMIXTURE
Neighbour Joining tree
VCF2Dis
Variant filtration
LD decay
PopLDdecay
MQ<40, QD<2,
FS>60 and 2<DP<50
Population differentiation
VCFtools
VCFtools
Nucleotide diversity
XP-CLR
Selective sweeps

### Slide 2
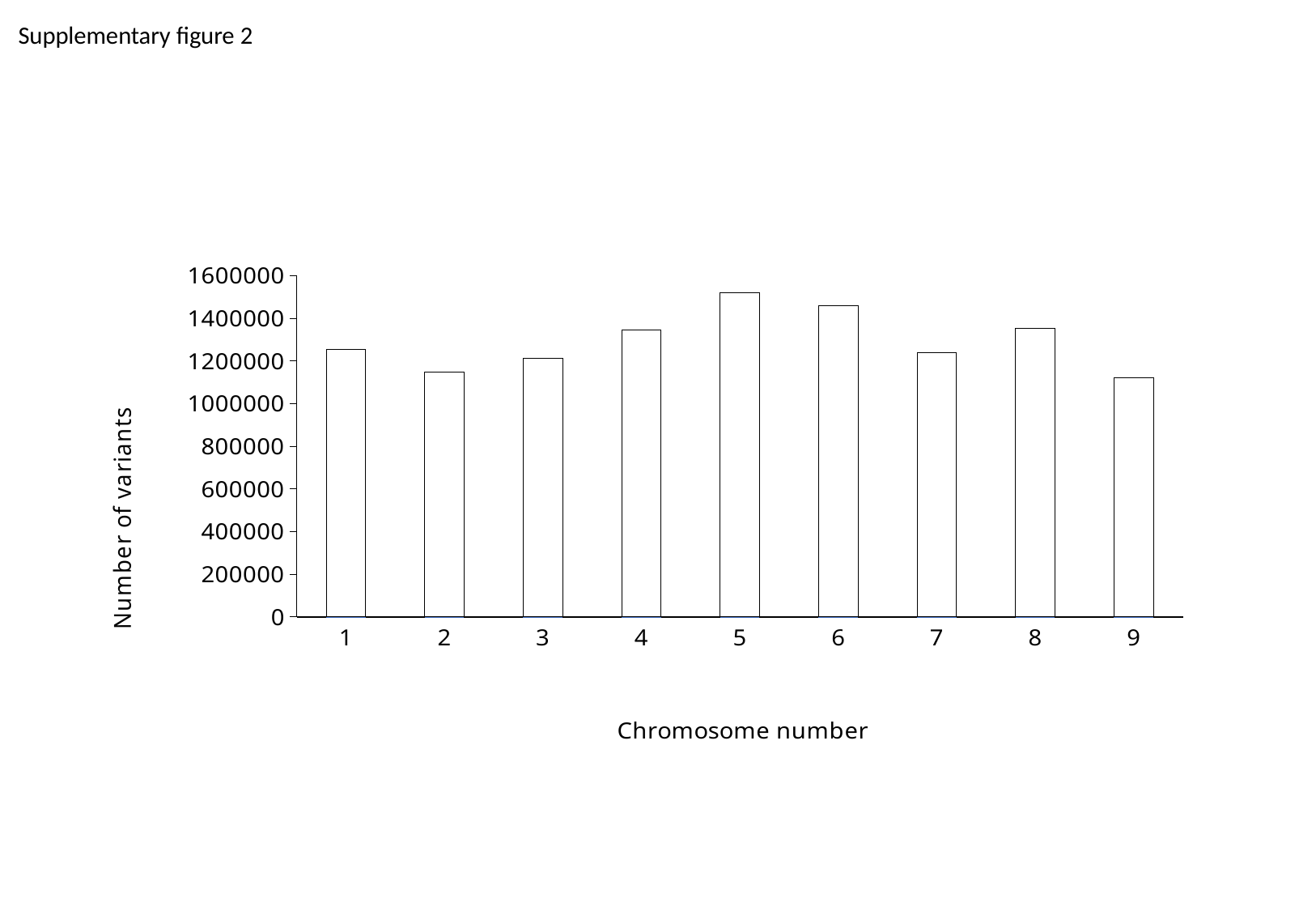

Supplementary figure 2
#### Chart
| Category |
|---|

### Slide 3
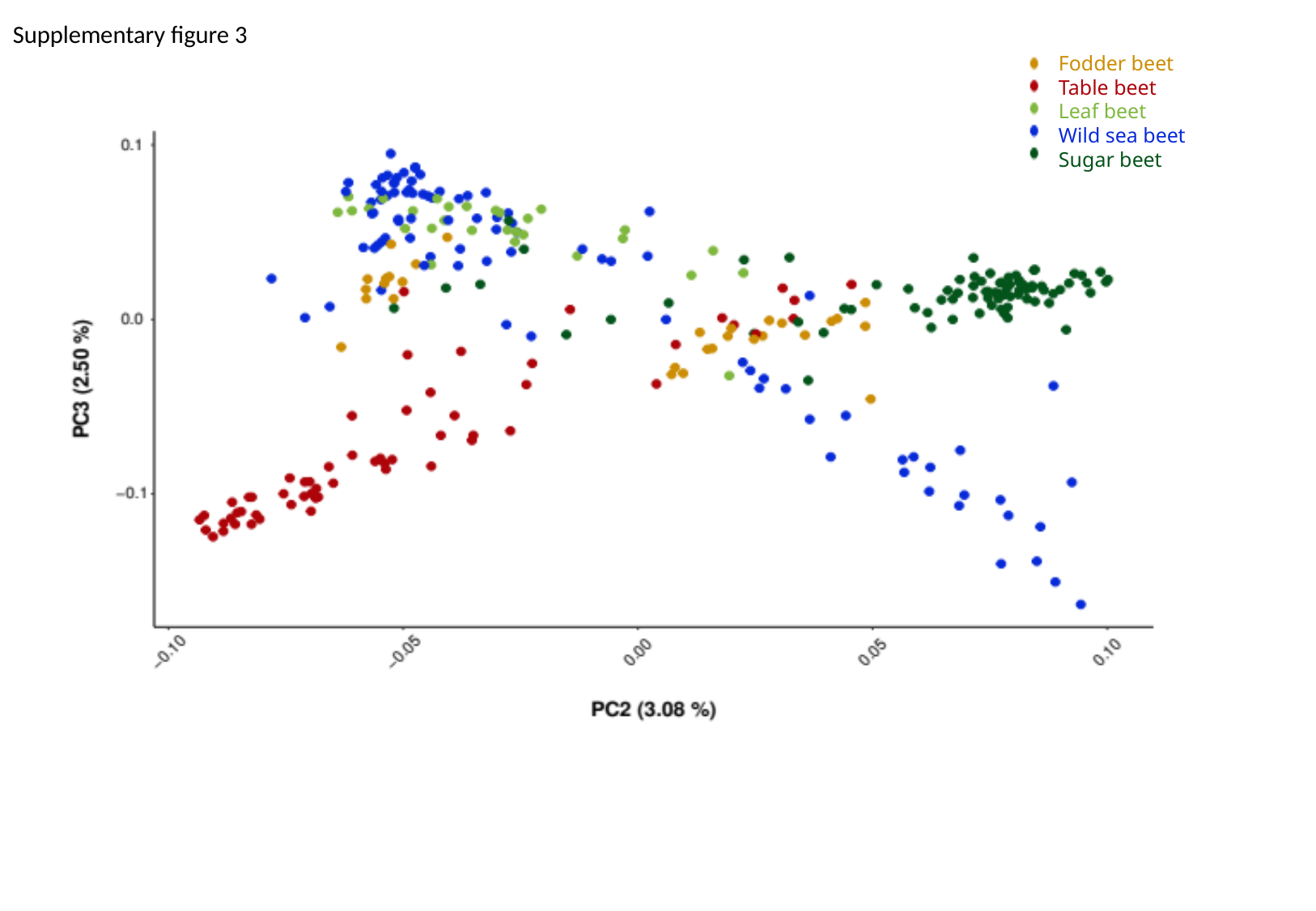

Supplementary figure 3
Fodder beet
Table beet
Leaf beet
Wild sea beet
Sugar beet

### Slide 4
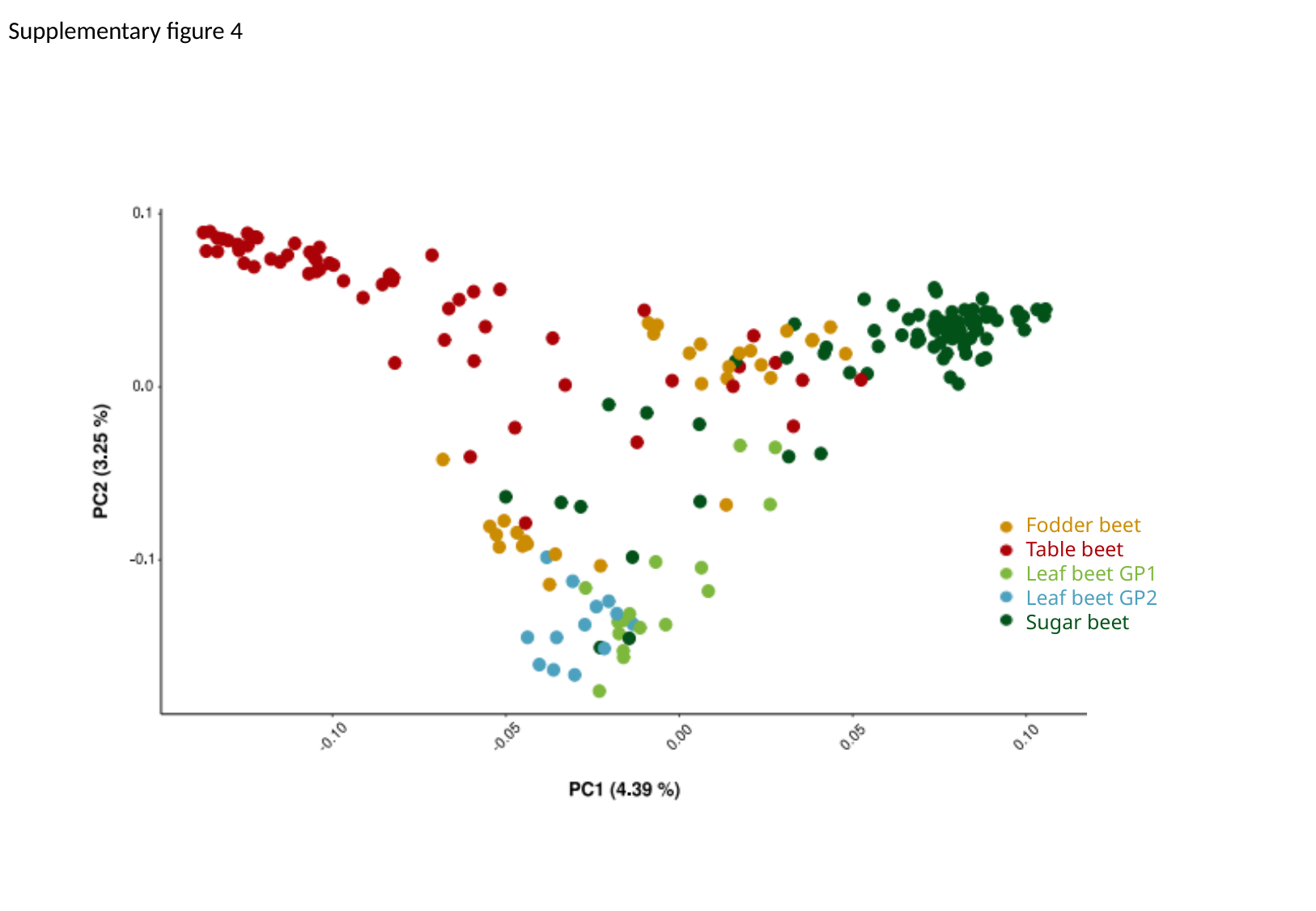

Supplementary figure 4
Fodder beet
Table beet
Leaf beet GP1
Leaf beet GP2
Sugar beet

### Slide 5
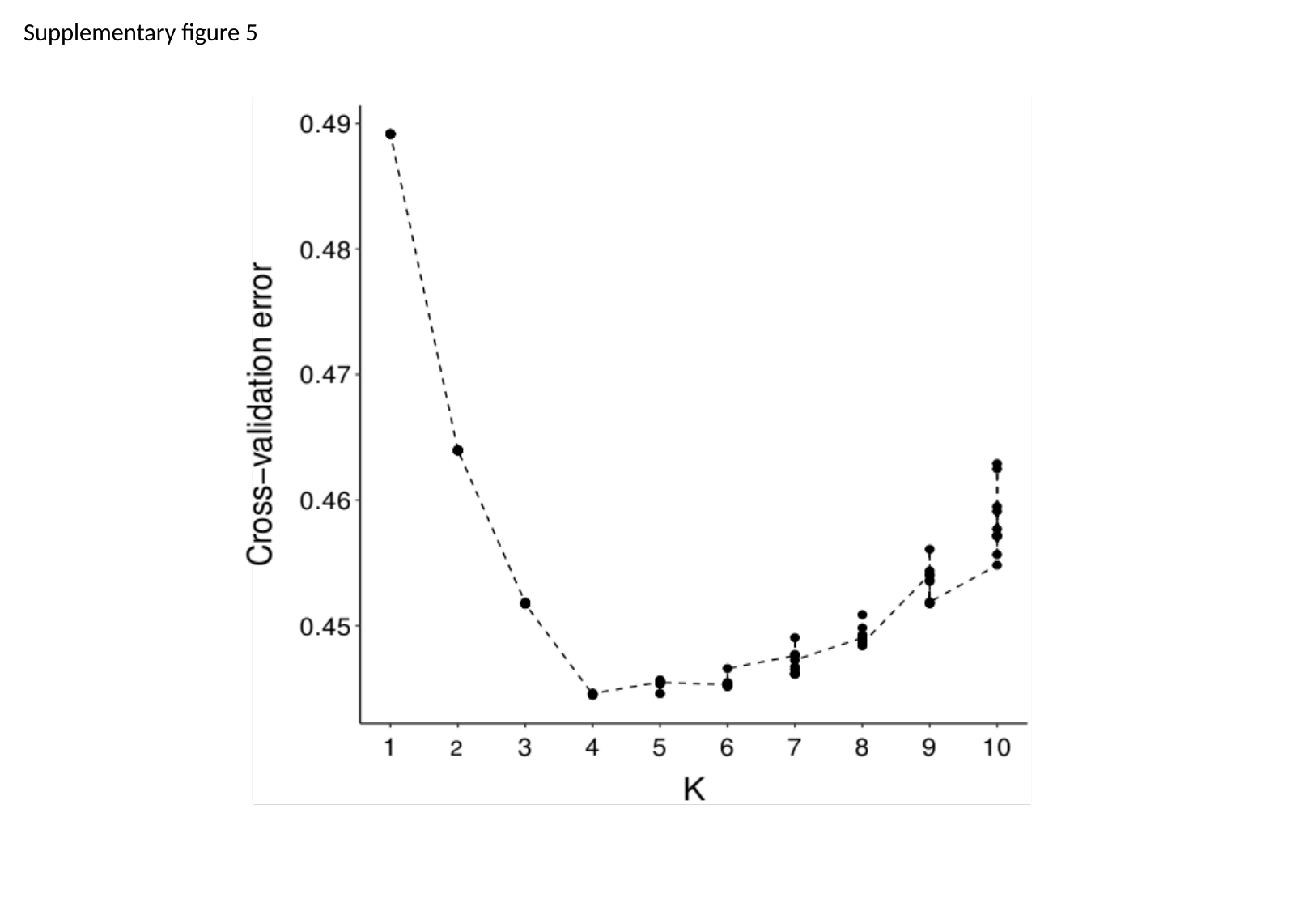

Supplementary figure 5

### Slide 6
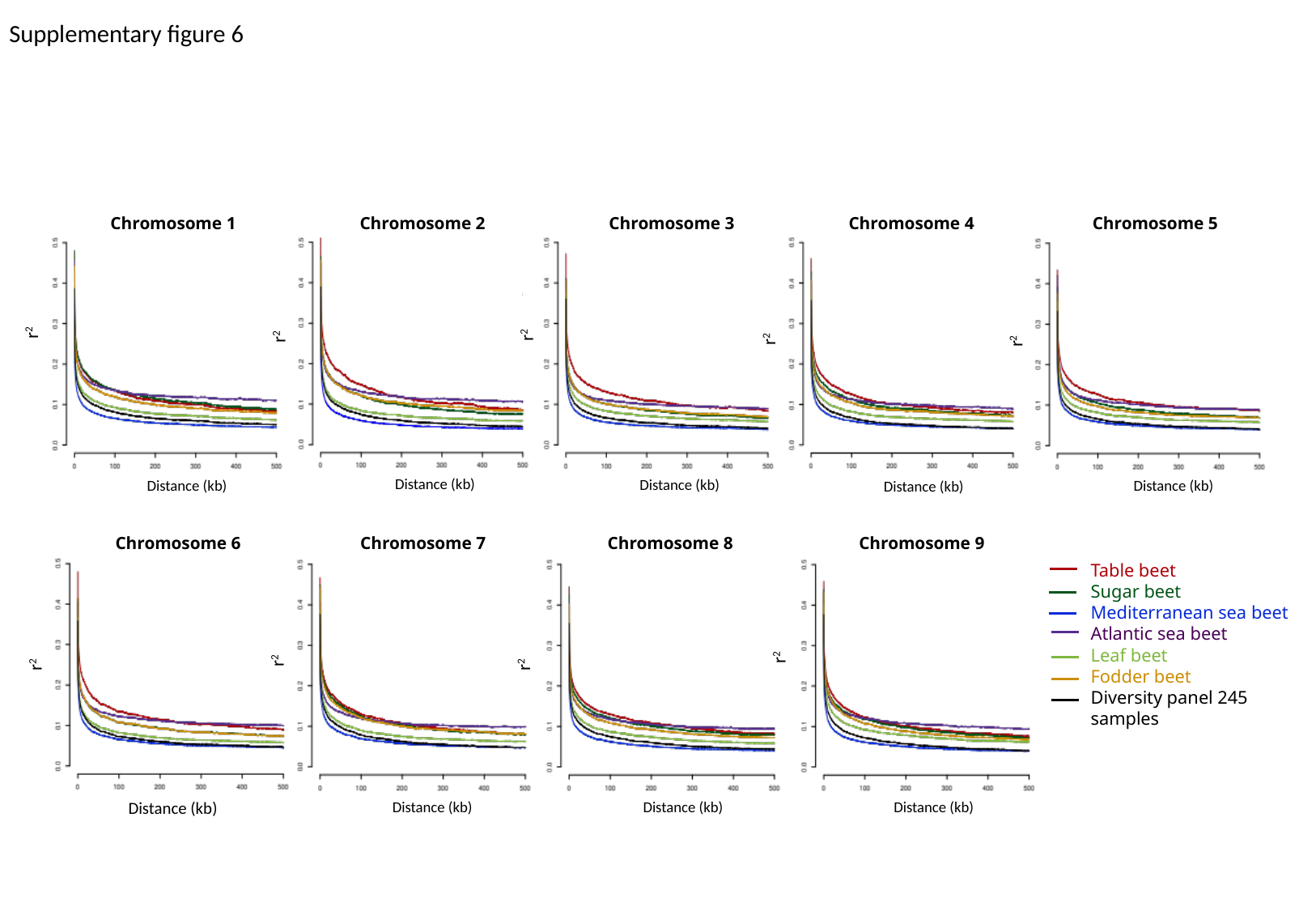

Supplementary figure 6
Chromosome 4
Chromosome 5
Chromosome 2
Chromosome 3
Chromosome 1
r^2^
r^2^
r^2^
r^2^
r^2^
Distance (kb)
Distance (kb)
Distance (kb)
Distance (kb)
Distance (kb)
Distance (kb)
Chromosome 6
Chromosome 7
Chromosome 8
Chromosome 9
Table beet
Sugar beet
Mediterranean sea beet
Atlantic sea beet
Leaf beet
Fodder beet
Diversity panel 245 samples
r^2^
r^2^
r^2^
r^2^
Distance (kb)
Distance (kb)
Distance (kb)
Distance (kb)

### Slide 7
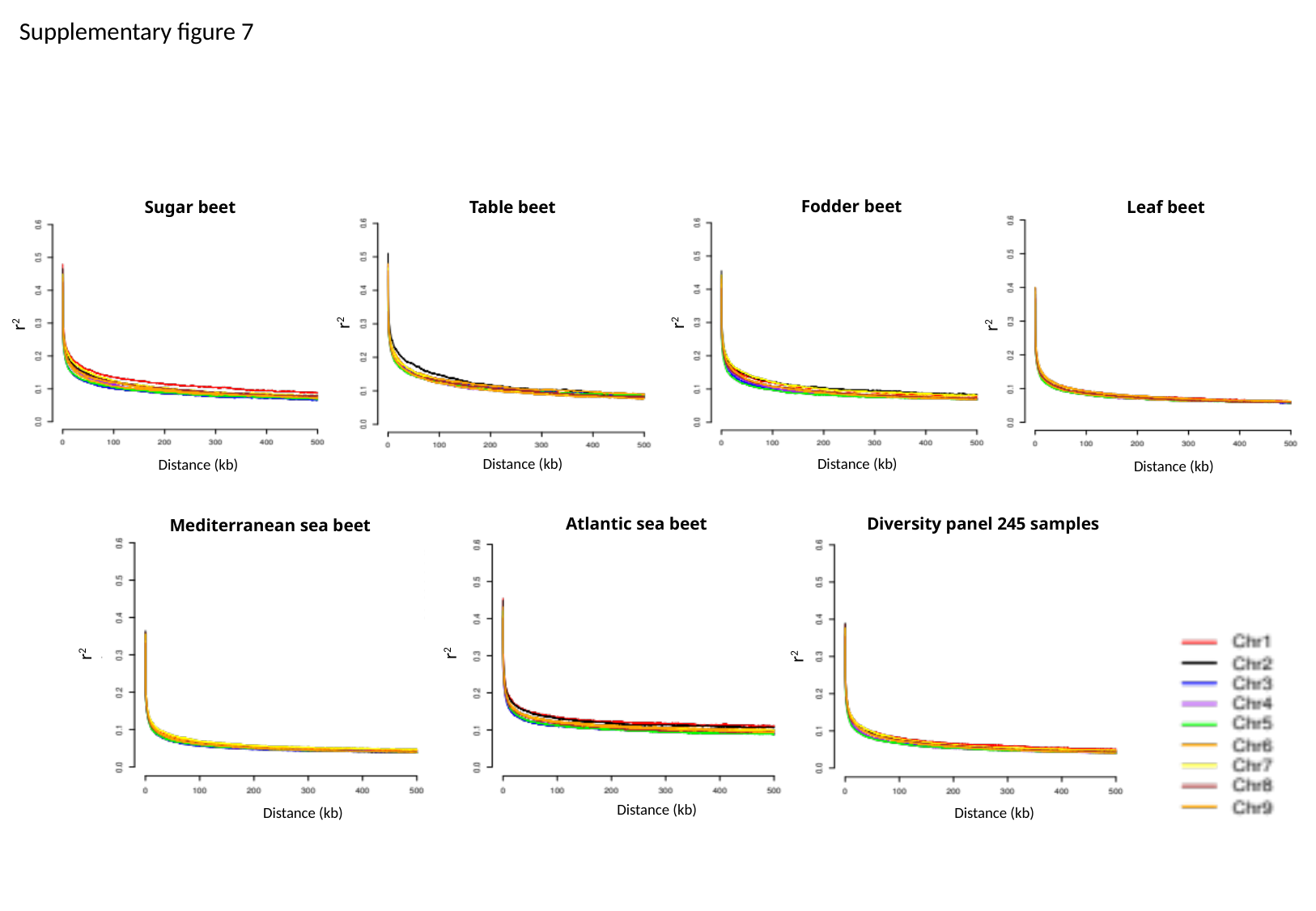

Supplementary figure 7
Fodder beet
Leaf beet
Table beet
Sugar beet
r^2^
r^2^
r^2^
r^2^
Distance (kb)
Distance (kb)
Distance (kb)
Distance (kb)
Atlantic sea beet
Diversity panel 245 samples
Mediterranean sea beet
r^2^
r^2^
r^2^
Distance (kb)
Distance (kb)
Distance (kb)

### Slide 8
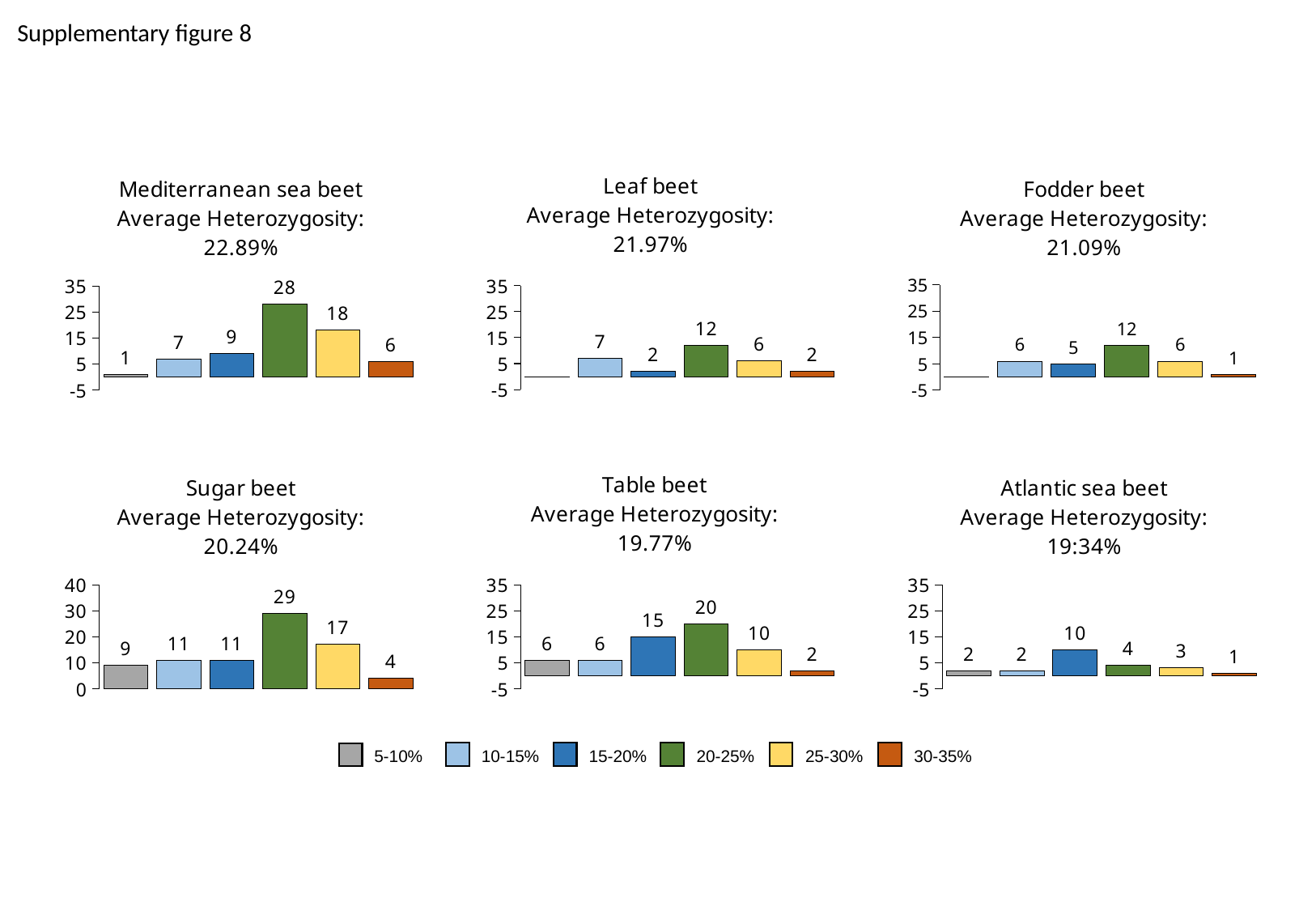

Supplementary figure 8
#### Chart: Leaf beet
Average Heterozygosity: 21.97%
| Category | |
|---|---|
| 5 | 0.0 |
| 10 | 7.0 |
| 15 | 2.0 |
| 20 | 12.0 |
| 25 | 6.0 |
| 30 | 2.0 |
#### Chart: Fodder beet
Average Heterozygosity: 21.09%
| Category | |
|---|---|
| 5 | 0.0 |
| 10 | 6.0 |
| 15 | 5.0 |
| 20 | 12.0 |
| 25 | 6.0 |
| 30 | 1.0 |
#### Chart: Mediterranean sea beet
Average Heterozygosity: 22.89%
| Category | |
|---|---|
| 5 | 1.0 |
| 10 | 7.0 |
| 15 | 9.0 |
| 20 | 28.0 |
| 25 | 18.0 |
| 30 | 6.0 |
#### Chart: Sugar beet
Average Heterozygosity: 20.24%
| Category | |
|---|---|
| 5 | 9.0 |
| 10 | 11.0 |
| 15 | 11.0 |
| 20 | 29.0 |
| 25 | 17.0 |
| 30 | 4.0 |
#### Chart: Table beet
Average Heterozygosity: 19.77%
| Category | |
|---|---|
| 5 | 6.0 |
| 10 | 6.0 |
| 15 | 15.0 |
| 20 | 20.0 |
| 25 | 10.0 |
| 30 | 2.0 |
#### Chart: Atlantic sea beet
Average Heterozygosity: 19:34%
| Category | |
|---|---|
| 5 | 2.0 |
| 10 | 2.0 |
| 15 | 10.0 |
| 20 | 4.0 |
| 25 | 3.0 |
| 30 | 1.0 |25-30%
30-35%
5-10%
10-15%
15-20%
20-25%

### Slide 9
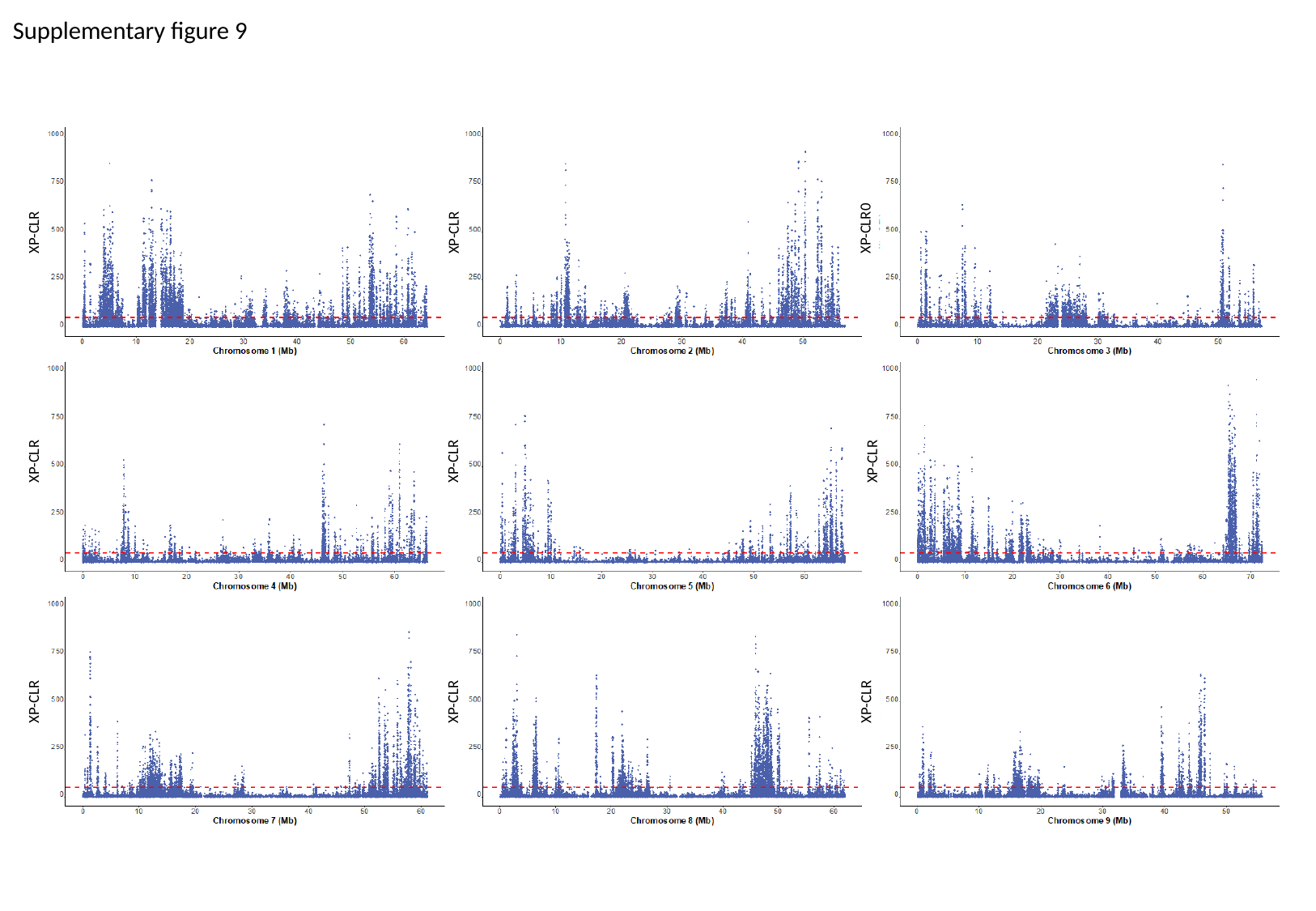

Supplementary figure 9
XP-CLR
XP-CLR0
XP-CLR
XP-CLR
XP-CLR
XP-CLR
XP-CLR
XP-CLR
XP-CLR

### Slide 10
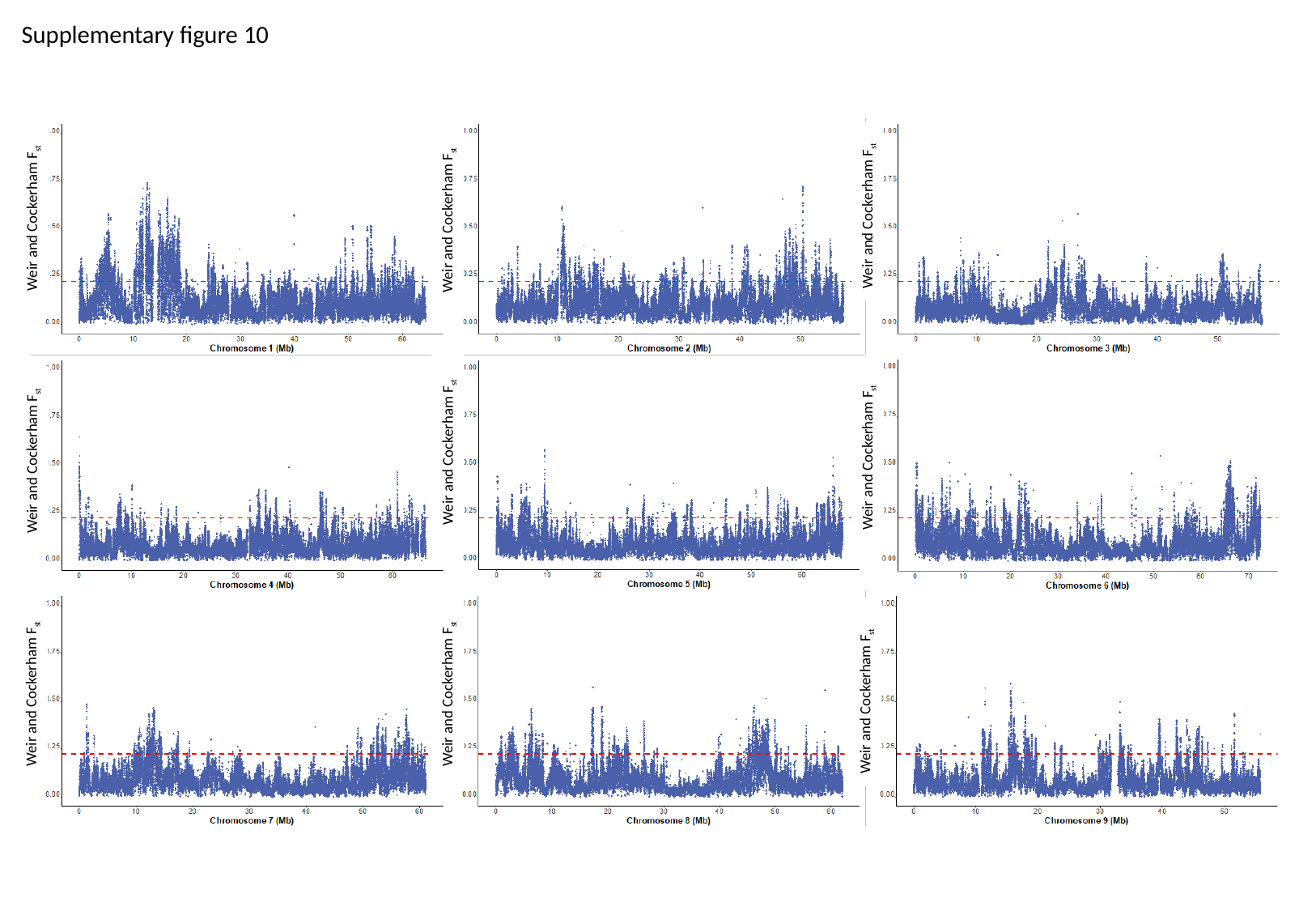

Supplementary figure 10
Weir and Cockerham F_st_
Weir and Cockerham F_st_
Weir and Cockerham F_st_
Weir and Cockerham F_st_
Weir and Cockerham F_st_
Weir and Cockerham F_st_
Weir and Cockerham F_st_
Weir and Cockerham F_st_
Weir and Cockerham F_st_

### Slide 11
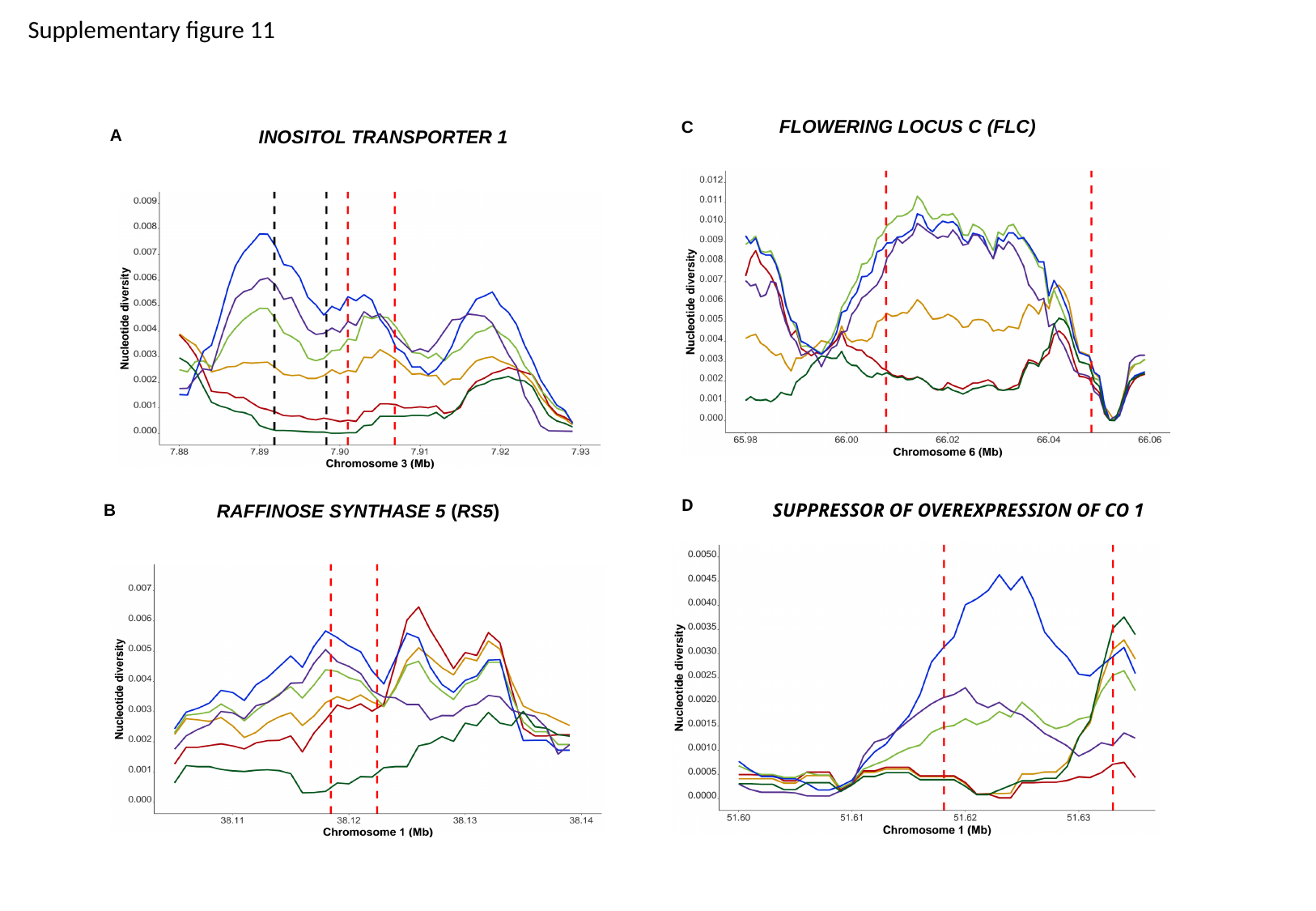

Supplementary figure 11
Flowering locus C (FLC)
C
A
Inositol transporter 1
D
SUPPRESSOR OF OVEREXPRESSION OF CO 1
B
Raffinose Synthase 5 (RS5)

### Slide 12
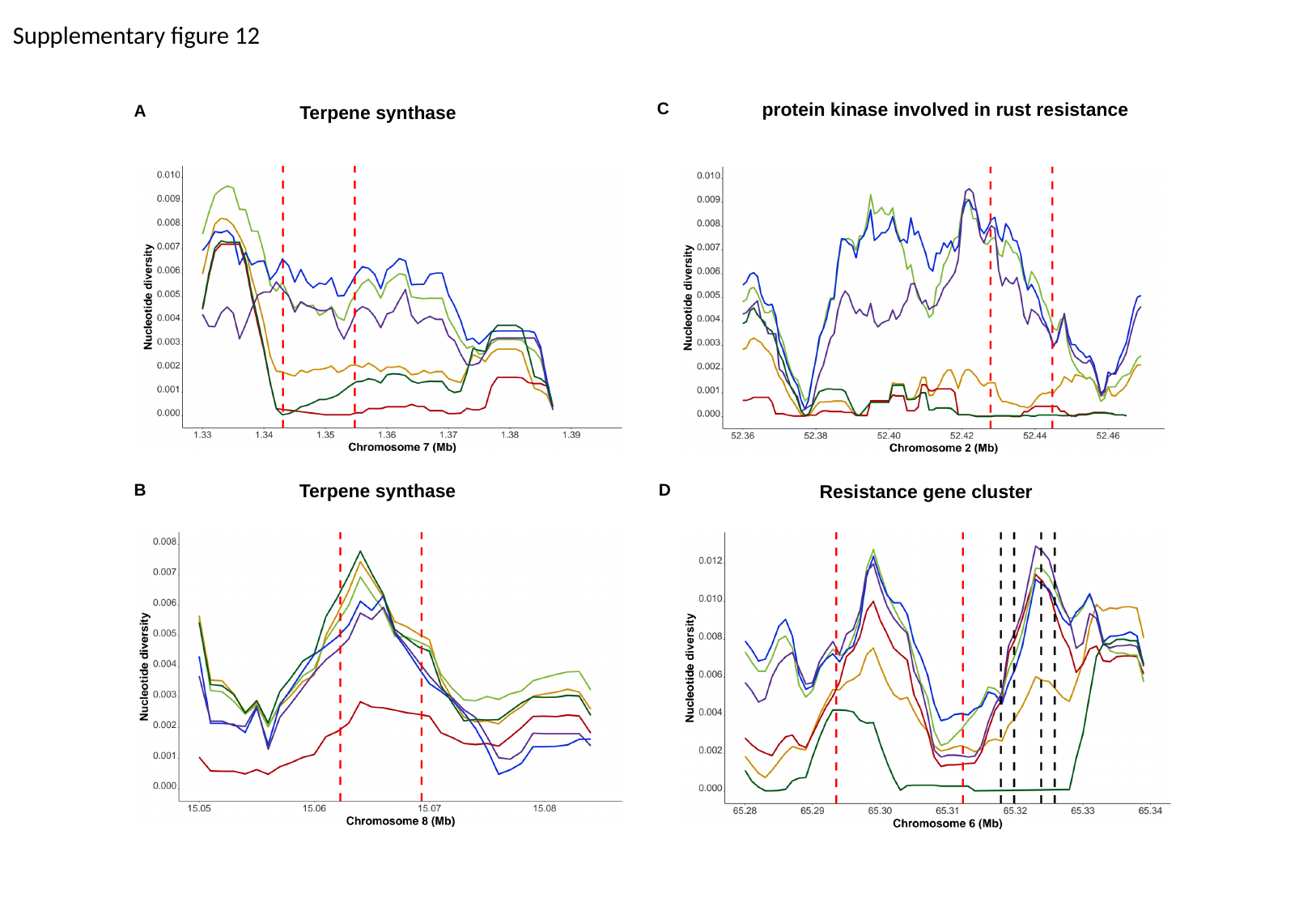

Supplementary figure 12
protein kinase involved in rust resistance
C
A
Terpene synthase
B
Terpene synthase
D
Resistance gene cluster

### Slide 13
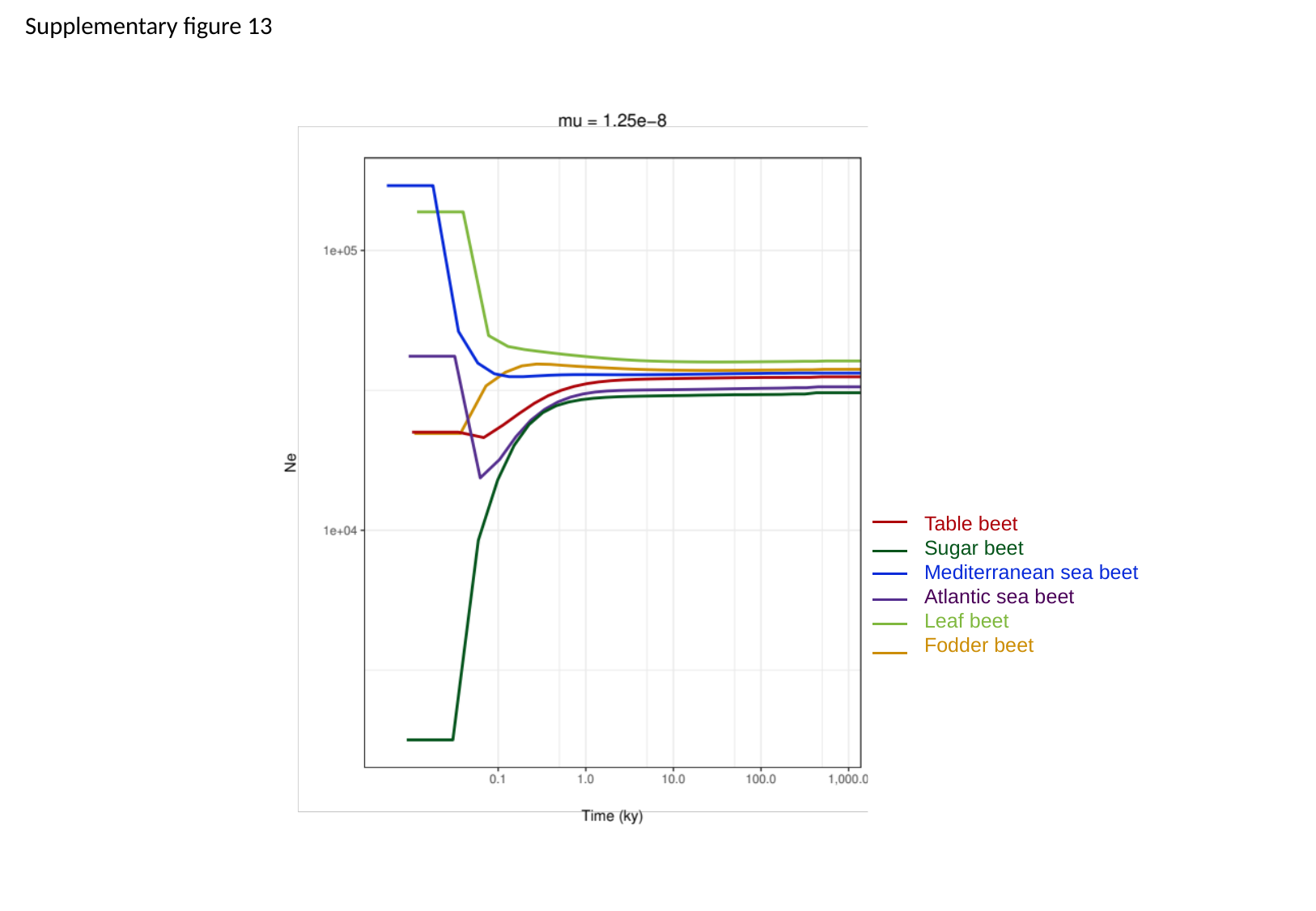

Supplementary figure 13
Table beet
Sugar beet
Mediterranean sea beet
Atlantic sea beet
Leaf beet
Fodder beet
